# Polymorphic structures of rapidly twisting 40-residue amyloid-β fibrils

**DOI:** 10.64898/2026.04.10.717728

**Authors:** Motahareh G. Larimi, Kent R. Thurber, Robert Tycko

## Abstract

Fibrils formed by 40- and 42-residue amyloid-β peptides (Aβ40 and Aβ42) are polymorphic, containing molecular structures that vary with growth conditions in ways that are not fully understood. Here we use cryogenic electron microscopy to characterize the structure of rapidly twisting Aβ40 fibrils, for which the distance between apparent width minima in electron microscope images (“cross-over distances”) is approximately 25 nm. From samples grown under a single set of growth conditions, we obtain high-resolution structures for three different rapidly twisting polymorphs. Although their cross-over distances are similar, the three rapidly twisting polymorphs differ in twist handedness, symmetry, molecular conformations, and intermolecular contacts. Two of the rapidly twisting polymorphs resemble slowly twisting Aβ40 polymorphs that have been described previously, including polymorphs extracted from brain tissue of Alzheimer’s disease patients or created by seeded growth from amyloid in brain tissue, but with shorter conformationally ordered segments and other specific conformational differences. These results contribute to our understanding of amyloid polymorphism, connections between morphology and molecular structure, and relationships between brain-derived and *in vitro*-grown fibrils.

## Introduction

Electron microscope images of amyloid fibrils formed by a single peptide or protein often show multiple coexisting morphologies (1–4). Although structural studies would be greatly simplified if all fibril morphologies contained the same molecular structure, solid state nuclear magnetic resonance (ssNMR) measurements on fibrils formed by amyloid-β peptides (3,5) and other polypeptides (6,7) in the first decade of the 21^st^ century showed that fibrils with different morphologies exhibit different sets of ssNMR signals and hence contain fundamentally different molecular structures. The molecular-level polymorphism of amyloid fibrils remains a subject of active research with the advent of modern cryogenic electron microscopy (cryo-EM) technology and its widespread application to amyloid structures (8–17).

Cryo-EM methods have been applied to both 40-residue and 42-residue amyloid-β (Aβ40 and Aβ42) fibrils, including fibrils extracted from human Alzheimer’s disease (AD) tissue (10–12,18) and transgenic mouse brain tissue (18,19), fibrils produced by seeded growth from AD brain tissue extracts (14,15,20,21), and fibrils grown entirely *in vitro* (8,13,15–17,22). Structural models developed from ssNMR data first showed that peptide conformations (also called “folds”) within the cross-β subunits (also called “protofilaments”) of Aβ40 and Aβ42 fibrils could be qualitatively different from one another (5,23–27). Cryo-EM studies have identified additional conformations and modes of interaction between subunits in both Aβ40 fibrils (10,12–20,22) and Aβ42 fibrils (11,19,21,28).

In the work described below, we characterized the conformational states and intermolecular interactions of Aβ40 in fibril polymorphs that have not been previously characterized at high resolution. We focused specifically on rapidly twisting Aβ40 fibrils, *i.e.*, fibrils for which the period of apparent width modulation in transmission electron microscope (TEM) and cryo-EM images, commonly called the “cross-over distance”, is approximately 25 nm. This choice was motivated by two considerations: (i) Cross-over distances in Aβ40 fibrils for which molecular models have been developed to date range from about 36 nm to over 100 nm. Since variations in cross-over distances may correlate with conformational variations, Aβ40 fibrils with cross-over distances less than 30 nm were likely candidates for novel molecular conformations; (ii) While S-shaped peptide conformations are common in Aβ42 fibrils (11,25–28) and have been observed in disease-associated mutant Aβ40 fibrils from transgenic mice and human familial AD brain tissue (18,19), wild-type Aβ40 fibril polymorphs with S-shaped peptide conformations have not been reported. Cross-over distances in brain-derived Aβ42 fibrils with S-shaped conformations are approximately 30 nm (11). It therefore seemed possible that Aβ40 conformations in rapidly twisting Aβ40 fibrils might resemble the S-shaped conformations in rapidly twisting Aβ42 fibrils. Although S-shaped conformations for Aβ40 were not identified in this work, several unexpected results were obtained: (i) The ensemble of rapidly twisting Aβ40 fibrils was found to include three distinct polymorphs with 24-32 nm cross-over distances. One additional polymorph with a closely related structure and with a 44 nm cross-over distance was also identified; (ii) Of the three main polymorphs, for which cryo-EM density maps with resolution better than 2.8 Å were obtained, two have right-handed twist and one has left-handed twist. Since all fibrils were grown *in vitro* (and simultaneously), this finding argues against the possibility that only brain-derived Aβ40 fibrils have right-handed twist (15); (iii) Although two of the polymorphs with high-resolution density maps contain two cross-β subunits that are related by quasi-2_1_ or C_2_ symmetry about the fibril growth direction, the third polymorph contains two inequivalent cross-β subunits. Coexistence of conformationally inequivalent subunits within the core of a single wild-type Aβ40 or Aβ42 fibril polymorph has not been reported previously; (iv) The structure within one rapidly twisting Aβ40 polymorph is nearly identical to the structure within slowly twisting, brain-seeded Aβ40 fibrils reported by Ghosh *et al*. (20), but with a longer disordered N-terminal segment. Comparison of rapidly twisting and slowly twisting polymorph structures indicates that large changes in cross-over distance can result from variations in the length of structurally ordered segments, together with subtle changes in molecular conformation and intermolecular contacts; (v) The structure within another rapidly twisting Aβ40 polymorph resembles structures reported for Aβ40 fibrils that were directly extracted from AD brain tissue (10,12) or amplified from tissue by seeded growth (14), but with a disordered N-terminal segment and different sidechain orientations towards the C-terminus in the rapidly twisting fibrils. These observations contribute to our understanding of the relationships among directly-extracted fibril polymorphs, polymorphs in brain-seeded samples, and fibril polymorphs that are prepared entirely *in vitro*.

## Results

### Preparation and imaging of rapidly twisting Aβ40 fibrils

Aβ40 fibrils were grown *in vitro* at 24° C or 37° C at various peptide concentrations, NaCl concentrations, and pH values, without stirring or agitation and without addition of pre-formed fibril seeds. Synthetic Aβ40 was used as the starting material. Fibril morphologies were assessed initially with TEM imaging (Fig. S1). A large population of rapidly twisting fibrils was found in samples grown at 37° C with 200 μM peptide concentration, 10 mM sodium phosphate buffer, pH 7.4, and no additional NaCl (Fig. S2). Cryo-EM images were then recorded with imaging parameters given in Table 1. Fig. 1A shows a representative cryo-EM image in which rapidly twisting fibrils are observed, along with slowly twisting and apparently untwisted fibrils. After fibril picking and reference-free two-dimensional (2D) classification in RELION software (29,30), three rapidly twisting fibril polymorphs were identified. Based on differences in symmetry discussed below, we refer to these polymorphs as RT-Aβ40(2_1_), RT-Aβ40(C_2_), and RT-Aβ40(C_1_) fibrils. Fig. 1B shows examples of 2D class average images of these three polymorphs, for which high-resolution three-dimensional (3D) density maps and molecular structural models could be obtained. Additional cryo-EM images and 2D class average images for all rapidly twisting polymorphs are shown in Figs. S3 and S4.

**Figure 1:**
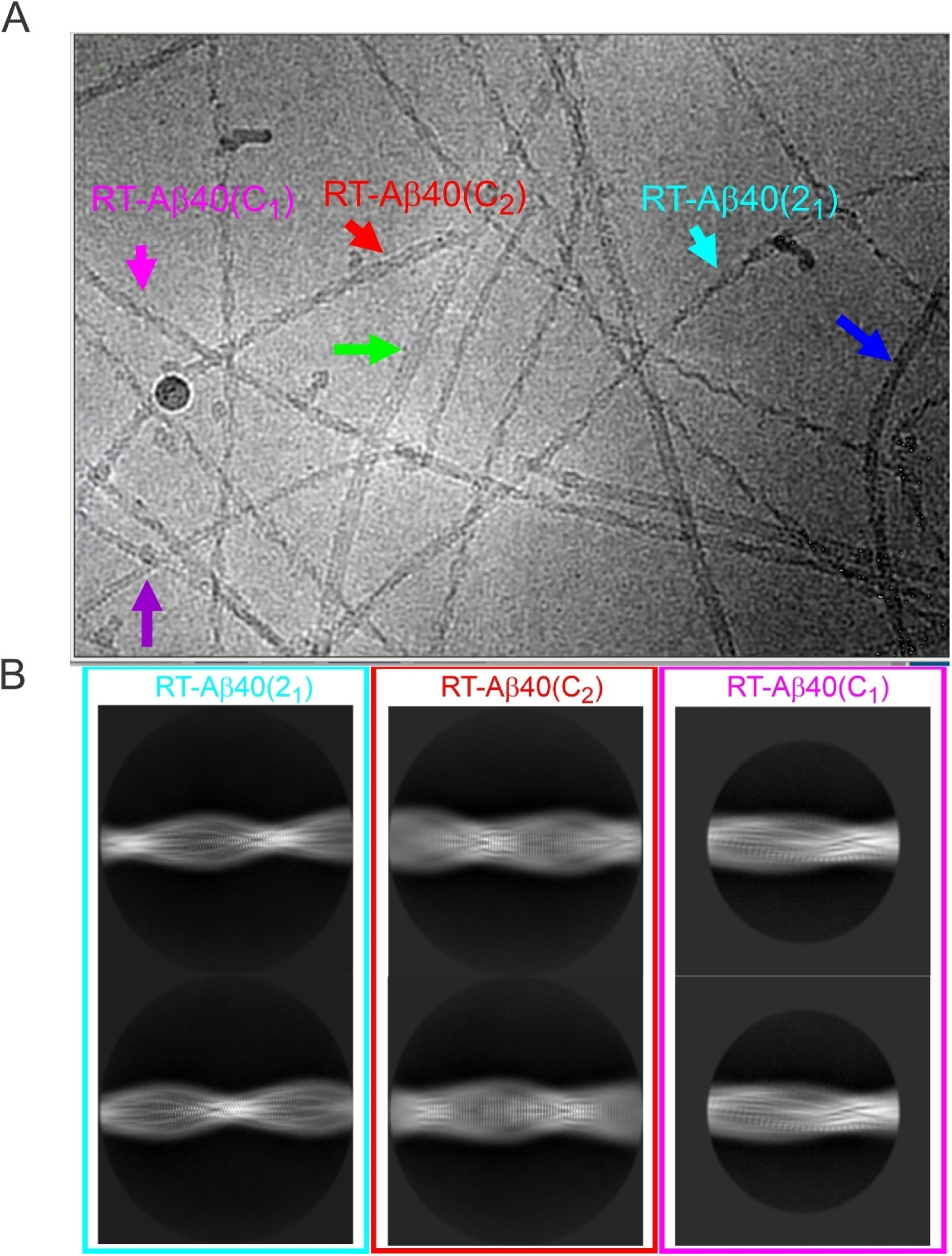
Polymorphic Aβ40 fibrils grown quiescently at 37° C in 10 mM sodium phosphate buffer, pH 7.4, with 200 μM Aβ40 concentration. (A) Representative cryo-EM image with cyan, red, and magenta arrows indicating examples of RT-Aβ40(2_1_), RT-Aβ40(C_2_), and RT-Aβ40(C_1_) fibrils, for which molecular structural models were developed in this work. Additional polymorphs are indicated by blue, green, and purple arrows. (B) Examples of 2D class average images of RT-Aβ40(2_1_), RT-Aβ40(C_2_), and RT-Aβ40(C_1_) fibrils.

**Table 1:**
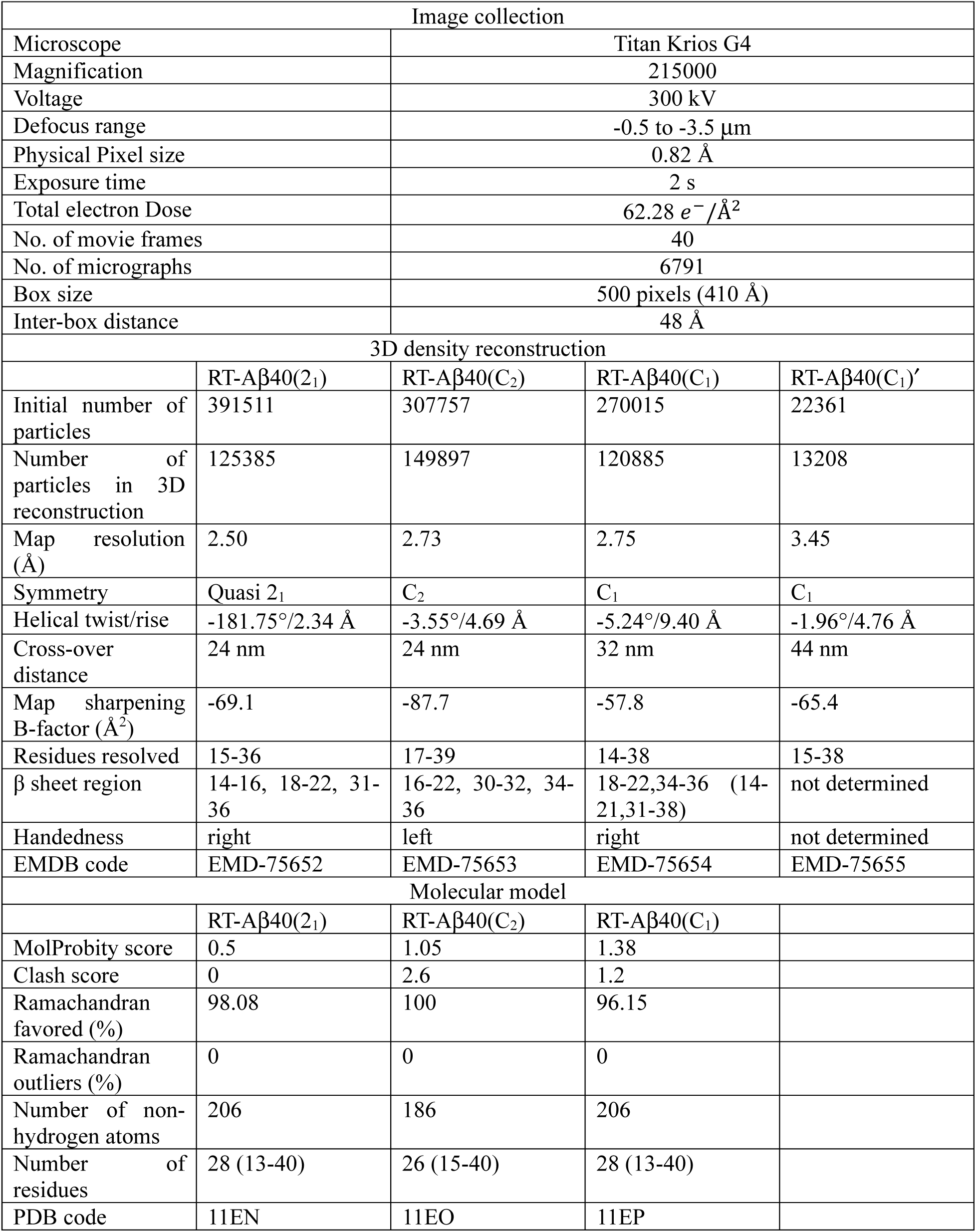
Parameters for cryo-EM image collection, 3D density reconstructions, and molecular models.

Based on 2D classification, sets of particles (*i.e.*, fibril segments) were prepared separately for each polymorph. Multiple rounds of three-dimensional (3D) classification and refinement were then performed to generate density maps, again using RELION software supplemented by custom scripts (20). Molecular models were created initially with Coot software (31). Initial models for RT-Aβ40(2_1_), RT-Aβ40(C_2_), and RT-Aβ40(C_1_) fibrils were then refined with Xplor-NIH calculations (32) as previously described (20,21). Relevant parameters for 3D classification, molecular model generation, and Xplor-NIH calculations are given in Table 1 and Table S1. This approach to molecular model generation results in bundles of structures from independent Xplor-NIH calculations, which provide estimates of the precision of atomic coordinates and torsion angles in the models (Fig. S5). Structure bundles for RT-Aβ40(2_1_), RT-Aβ40(C_2_), and RT-Aβ40(C_1_) fibrils were deposited in the Protein Data Bank (PDB) with PDB codes 11EN, 11EO, and 11EP, respectively.

In 3D classification of the RT-Aβ40(C_1_) particles with four allowed classes, a related polymorph which we call RT-Aβ40(C_1_)′ was identified. Particles assigned to the RT-Aβ40(C_1_)′ class were subsequently analyzed separately. 3D classification of RT-Aβ40(2_1_) and RT-Aβ40(C_2_) particles with four allowed classes produced four indistinguishable density maps, indicating that these sets of particles were essentially monomorphic.

### Structure of RT-Aβ40(2_1_) fibrils

Figs. 2A and 2B show the 3D density map for RT-Aβ40(2_1_) fibrils, for which a final resolution of 2.50 Å was obtained by imposing quasi-2_1_ symmetry in addition to overall helical symmetry. Density maps calculated without additional symmetry or with C_2_ symmetry had somewhat lower final resolutions (Fig. S6A). Fig. 2C shows the molecular model in a cross-sectional view of the density, which includes two symmetry-related cross-β subunits. The conformationally ordered core structure is formed by residues 15-36. Residues 1-14 and 37-40 are apparently disordered or adopt multiple conformations, contributing to a diffuse halo of density around the ordered core (Fig. S7A). Expanded views in Figs. 2D and 2E show inter-subunit hydrophobic contacts, involving sidechains of L17, F19, and V24 from one subunit and sidechains of I32, L34, and V36 from the other subunit, as well as “steric zipper” packing (33) between residues 25-27 of the two subunits. Sidechains of N27 in each subunit appear likely to engage in “polar zipper” interactions (34), and sidechains of D23 and K28 engage in salt bridge interactions (35).

**Figure 2:**
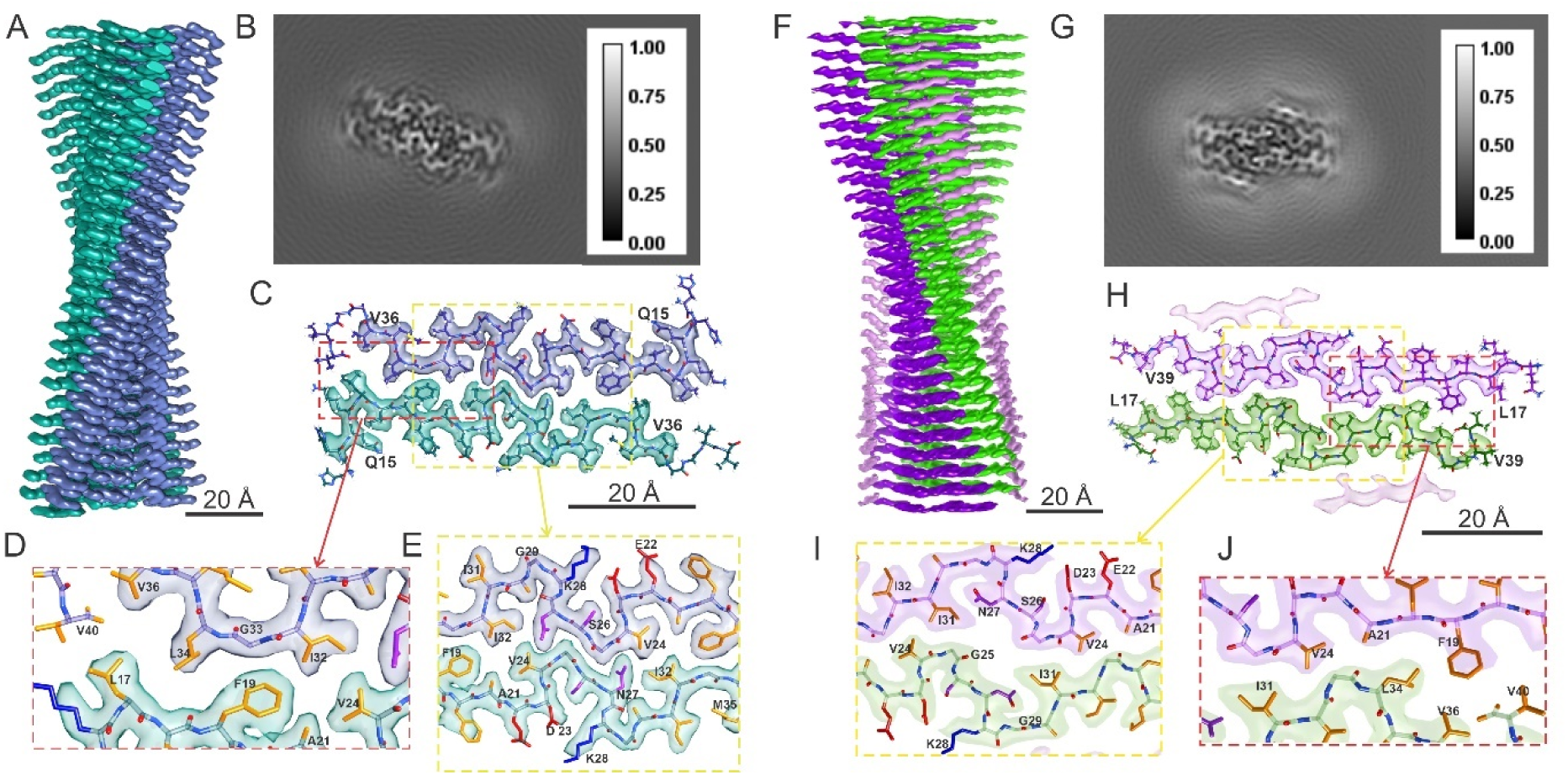
Cryo-EM density maps and molecular models for RT-Aβ40(2_1_) and RT-Aβ40(C_2_) fibrils. (A) Side view of a surface rendering of the RT-Aβ40(2_1_) density map, with right-handed twist, quasi-2_1_ symmetry, and 2.50 Å resolution. The two cross-β subunits are colored blue and green. (B) Cross-sectional view of the RT-Aβ40(2_1_) density map, summed over 7 pixels along fibril growth axis. (C) Cross-sectional view of the molecular model for RT-Aβ40(2_1_) fibrils superimposed on one layer of the density map. (D,E) Expanded views showing hydrophobic sidechain contacts between the two subunits and a salt bridge interaction between sidechains of K28 and D23. Hydrophobic, basic, acidic and polar side chain atoms are shown in yellow, blue, red and purple, respectively. (F-J) Same as panels A-E, but for the RT-Aβ40(C_2_) polymorph, with left-handed twist, C_2_ symmetry, and 2.73 Å resolution. Cross-β subunits are colored purple and light green.

As is well known, the handedness of helical symmetry cannot be determined directly from cryo-EM images of amyloid fibrils but can be determined by comparison of left-handed and right-handed density maps with their corresponding molecular models when the resolution is better than about 3 Å. The model in Fig. 2C is based on the density map with right-handed twist in Fig. 2A. Right-handed twist is supported by the alignment of backbone carbonyl oxygens in β-strand residues 18-21 with bulges in the density (Fig. S8A). In contrast, the molecular model for a density map with left-handed twist shows backbone carbonyl oxygens in residues 18-21 that do not align with bulges in the density.

### Structure of RT-Aβ40(C_2_) fibrils

Figs. 2F and 2G show the 3D density map for RT-Aβ40(C_2_) fibrils, for which a final resolution of 2.73 Å was obtained by imposing C_2_ symmetry (Fig. S6B). Comparison of left-handed and right-handed density maps with their corresponding molecular models favors a left-handed helical twist (Fig. S8B). Fig. 2H shows the molecular model in a cross-sectional view of the density. The conformationally ordered segment in this model includes residues 17-39. Expanded views in Figs. 2I and 2J show inter-subunit hydrophobic contacts in RT-Aβ40(C_2_) fibrils, which involve sidechains of F19, A21, and V24 from one subunit and sidechains of I31, L34, and V36 from the other subunit. Inter-subunit interactions are clearly different in RT-Aβ40(C_2_) and RT-Aβ40(2_1_) fibrils. For example, the RT-Aβ40(C_2_) model includes inter-subunit F19-L34, F19-V36, and V24-I31 contacts, whereas the RT-Aβ40(2_1_) model includes L17-L34, F19-I32, and V24-I32 contacts. The intra-subunit configurations of residues 22-28 are nearly the same in the two rapidly twisting polymorphs, apart from the conformation and contacts of N27 sidechains.

In addition to the two central cross-β subunits related by C_2_ symmetry that comprise the fibril core, the density map in Fig. 2H includes additional outer layers with density nearly equal to that of the central layers (Fig. S7B). These outer layers may represent partially disordered β-sheets, possibly formed by Aβ40 molecules with hairpin-like conformations as suggested in our previous study of brain-seeded Aβ40 fibrils (20). The outer layers cover hydrophobic sidechains of A30, I32, and M35 that would otherwise be exposed to solvent on the exterior of RT-Aβ40(C_2_) fibrils. The absence of similar outer layers in the density map for RT-Aβ40(2_1_) fibrils may be attributed to the different conformation of residues 30-35. In RT-Aβ40(2_1_) fibrils, sidechains of A30 and I32 are contained within the core, rather than being solvent-exposed. Although sidechains of I31 and M35 are exposed in RT-Aβ40(2_1_) fibrils, the effective hydrophobic surface area may be less than in RT-Aβ40(C_2_) fibrils. It is worth noting that Aβ40 fibrils grown in the presence of lipid vesicles by Frieg *et al*. (13) (PDB code 8OVK) had a core structure that closely resembles that of our RT-Aβ40(C_2_) fibrils, but with lipids covering the solvent-exposed hydrophobic surfaces.

### Structure of RT-Aβ40(C_1_) and RT-Aβ40(C_1_)′ fibrils

Figs. 3A and 3B show the 3D density map for RT-Aβ40(C_1_) fibrils with 2.75 Å resolution (Fig. S6C). Since the density map clearly shows two inequivalent cross-β subunits in the helical repeat unit (*i.e.*, C_1_ symmetry), calculations with pseudo-2_1_ or C_2_ symmetry imposed on the repeat unit were not performed. Comparison of molecular models with left-handed and right-handed densities supports a right-handed twist for RT-Aβ40(C_1_) fibrils (Fig. S8C).

**Figure 3:**
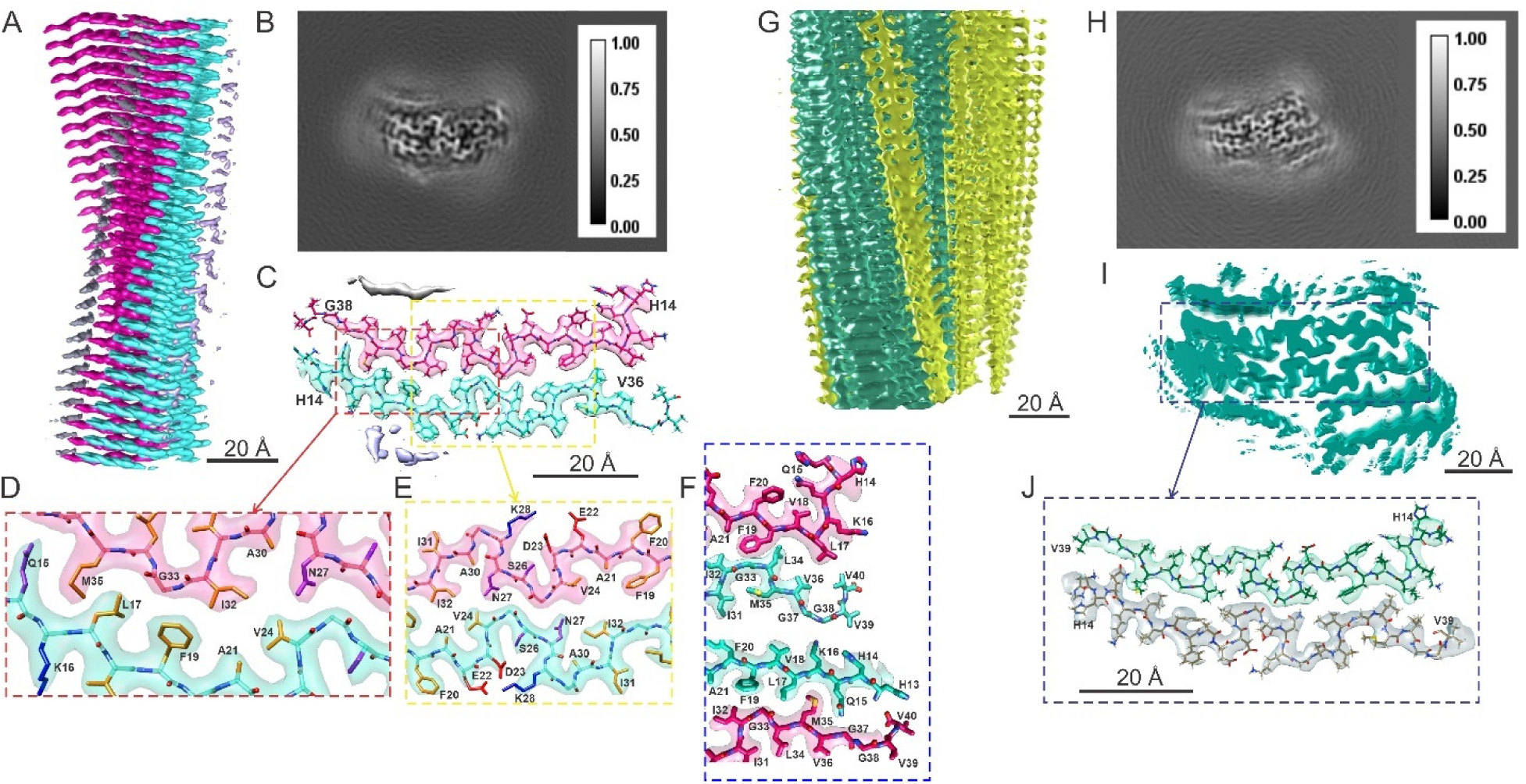
Cryo-EM density maps and molecular models for RT-Aβ40(C_1_) and RT-Aβ40(C_1_)′ fibrils. (A) Side view of a surface rendering of the RT-Aβ40(C_1_) density map, with right-handed twist and 2.75 Å resolution. “Upper” and “lower” cross-β subunits are colored magenta and cyan. (B) Cross-sectional view of the RT-Aβ40(C_1_) density map summed over 7 pixels along fibril growth axis. (C) Cross-sectional view of molecular model for RT-Aβ40(C_1_) fibrils superimposed on one layer of the density map with the molecular model. (D,E) Expanded views showing hydrophobic sidechain contacts between the two subunits and a salt bridge interaction between K28 and D23. Hydrophobic, basic, acidic and polar side chain atoms are shown in yellow, blue, red and purple, respectively. (F) Comparison of inter-subunit contacts between residues 13-19 of the upper subunit and residues 33-39 of the lower subunit or between residues 13-19 of the lower subunit and residues 33-39 of the upper subunit. Multiple differences in conformations and contacts illustrate the structural inequivalence of the two subunits in RT-Aβ40(C_1_) fibrils. (G,H) Same as panels A and B, but for the RT-Aβ40(C_1_)′ polymorph. (I) Surface rendering of a cross-sectional view of the RT-Aβ40(C_1_)′ density map, showing a structurally ordered central region that closely resembles the RT-Aβ40(C_1_) density map, with additional partially ordered regions above and below the upper and lower subunits. (J) Molecular model in the central region of the RT-Aβ40(C_1_)′ model, based on the molecular model for RT-Aβ40(C_1_) fibrils.

The molecular model in Fig. 3C shows that residues 14-37 in the “lower” (cyan) subunit and residues 15-37 in the “upper” (pink) subunit of RT-Aβ40(C_1_) fibrils are conformationally ordered. Expanded views in Figs. 3D and 3E show inter-subunit hydrophobic contacts between sidechains of L17, F19, A21, and V24 from the lower subunit and sidechains of I32 and M35 from the upper subunit, as well as hydrophobic contacts between sidechains of L17, F19, A21, and V24 from the upper subunit and sidechains of I32, L34, and V36 from the lower subunit. Conformations of residues 22-28 in both subunits of RT-Aβ40(C_1_) fibrils resemble the corresponding conformations in both RT-Aβ40(2_1_) and RT-Aβ40(C_2_) fibrils, with the orientations of N27 sidechains in RT-Aβ40(C_1_) fibrils being more similar to the orientation in RT-Aβ40(2_1_) fibrils.

Details of the inequivalence between subunits in RT-Aβ40(C_1_) fibrils are particularly obvious in Fig. 3F. Among other differences, M35 and I31 sidechains in the lower (cyan) subunit interact with one another, while M35 sidechains of the upper (pink) subunit interact with L17 sidechains of the lower subunit. Conformations of L17, F19, and F20 sidechains are different in the two subunits. In the lower subunit, residues 14-22 form a continues β-strand, while only residues 17-22 form a β-strand in the upper subunit.

In addition to the two central subunits in the density map for RT-Aβ40(C_1_) fibrils, weak layers of density occur above I31, L34, and V36 of the upper subunit and below V18 and F20 of the lower subunit (Figs 3B, 3C, and S7C), attributable to a low population of disordered Aβ40 molecules that adhere to exposed hydrophobic surfaces. The asymmetry of these weak density layers can be attributed to differences in the identities and conformations of solvent-exposed sidechains (I31, L34, and V36 from the upper subunit versus I31 and M35 from the lower subunit; F20 from the upper subunit versus V18 and F20 from the lower subunit) that create differences in hydrophobic surface areas.

A separate class of fibrils with the same core structure as RT-Aβ40(C_1_) fibrils, but with stronger density in additional layers (Fig. S7D) and a longer cross-over distance (Table 1), was also identified in the same set of cryo-EM images. Figs. 3G, 3H, and 3I show the 3D density map for these fibrils, which we call RT-Aβ40(C_1_)′ fibrils. Although the density map is depicted with left-handed twist, the lower resolution of this map (3.45 Å, Fig. S6D) precluded a definite determination of handedness. Fig. 3J shows that a molecular model that is nearly identical to the model for RT-Aβ40(C_1_) fibrils fits the central RT-Aβ40(C_1_)′ fibril density adequately.

### Comparisons among rapidly twisting Aβ40 fibrils

Although RT-Aβ40(2_1_), RT-Aβ40(C_2_), and RT-Aβ40(C_1_) fibrils have similar cross-over distances, their symmetries and the Aβ40 conformations in their cores are clearly different. The twist handedness of RT-Aβ40(C_2_) fibrils also differs from that of RT-Aβ40(2_1_) and RT-Aβ40(C_1_) fibrils. Thus, although the cross-over distance has been used to distinguish among fibril polymorphs in early studies (1,4), this parameter alone is not sufficient for unique identification.

If we define β-strand segments to be segments consisting of at least three consecutive residues whose backbone amide and carbonyl groups form hydrogen bonds to neighboring peptide chains on alternating sides, the β-strand segments in RT-Aβ40(2_1_) fibrils are residues 14-16, 18-22, and 31-36 (Fig. S9A). In RT-Aβ40(C_2_) fibrils, the β-strand segments are residues 16-22, 30-32, and 34-36 (Fig. S9B). In RT-Aβ40(C_1_) fibrils, β-strands are residues 18-22 and 34-36 in upper subunits and residues 14-21 and 31-36 in lower subunits (Fig. S9C). Thus, the β-strands in all three polymorphs are localized within residues 14-22 and 30-36, but with variations in their lengths and continuity.

Differences in β-strand segments among the three polymorphs are accompanied by differences in the identities of residues with backbone torsion angles that deviate from typical β-strand values (*i.e.*, φ ≍ −130° ± 30°, ψ ≍ 130° ± 30°). For example, the φ angles of L17 deviate strongly from β-strand values in RT-Aβ40(2_1_) fibrils and in the upper subunit of RT-Aβ40(C_1_) fibrils, but not in RT-Aβ40(C_2_) fibrils or in the lower subunit of RT-Aβ40(C_1_) fibrils (Fig. S10A). The φ and ψ angles of G33 deviate strongly from β-strand values in all polymorphs. However, in RT-Aβ40(2_1_) fibrils and in the lower subunit of RT-Aβ40(C_1_) fibrils, the φ and ψ angles of G33 differ from β-strand values only by a change in sign (Fig. S10B), allowing G33 to be included in continuous β-strands. The change in sign of these torsion angles accounts for the pronounced nonlinearity of the β-strands that include G33, as shown in Figs. 2D and 3C.

In all three polymorphs, residues 23-29 form similar turn-like configurations that bring sidechains of D23 and K28 into proximity. These configurations are determined by non-β-strand values for φ and/or ψ at D23, G25, N27, and K28. The precise values vary among the three polymorphs (Fig. S10B).

### Comparisons between RT-Aβ40(2_1_) fibrils and previously described polymorphs

The symmetry axis of RT-Aβ40(2_1_) fibrils is centered between S26 residues of the two cross-β subunits, with residues 24-27 participating in close inter-subunit contacts (Fig. 2E). Fig. 4A compares the conformation of Aβ40 in RT-Aβ40(2_1_) fibrils with conformations in three previously reported fibril polymorphs with similar inter-subunit contacts (12,14,15) (PDB codes 8FF2, 8QN6, and 8OT4) and in two additional polymorphs with related structures (10,17) (PDB codes 9IIO and 6SHS). Cross-over distances, twist handedness, and symmetries are given in Table 2. An obvious difference between RT-Aβ40(2_1_) fibrils and the other polymorphs in Fig. 4A is the absence of conformational order in residues 1-13 of RT-Aβ40(2_1_) fibrils (Figs. 2C and S7A). The shorter ordered segment in RT-Aβ40(2_1_) fibrils is one factor that may account for the more rapid twist. In addition, while the backbone and sidechain conformations of residues 21-32 are nearly identical in RT-Aβ40(2_1_) fibrils and in polymorphs represented by PDB codes 8FF2, 8QN6, and 8OT4, conformational differences in residues 33-40 and 14-20 lead to distinct sidechain directions and orientations and distinct contacts between cross-β subunits. Differences between RT-Aβ40(2_1_) fibrils and polymorphs represented by PDB codes 9IIO and 6SHS are more pronounced, as shown by superpositions in Fig. 4B.

**Figure 4:**
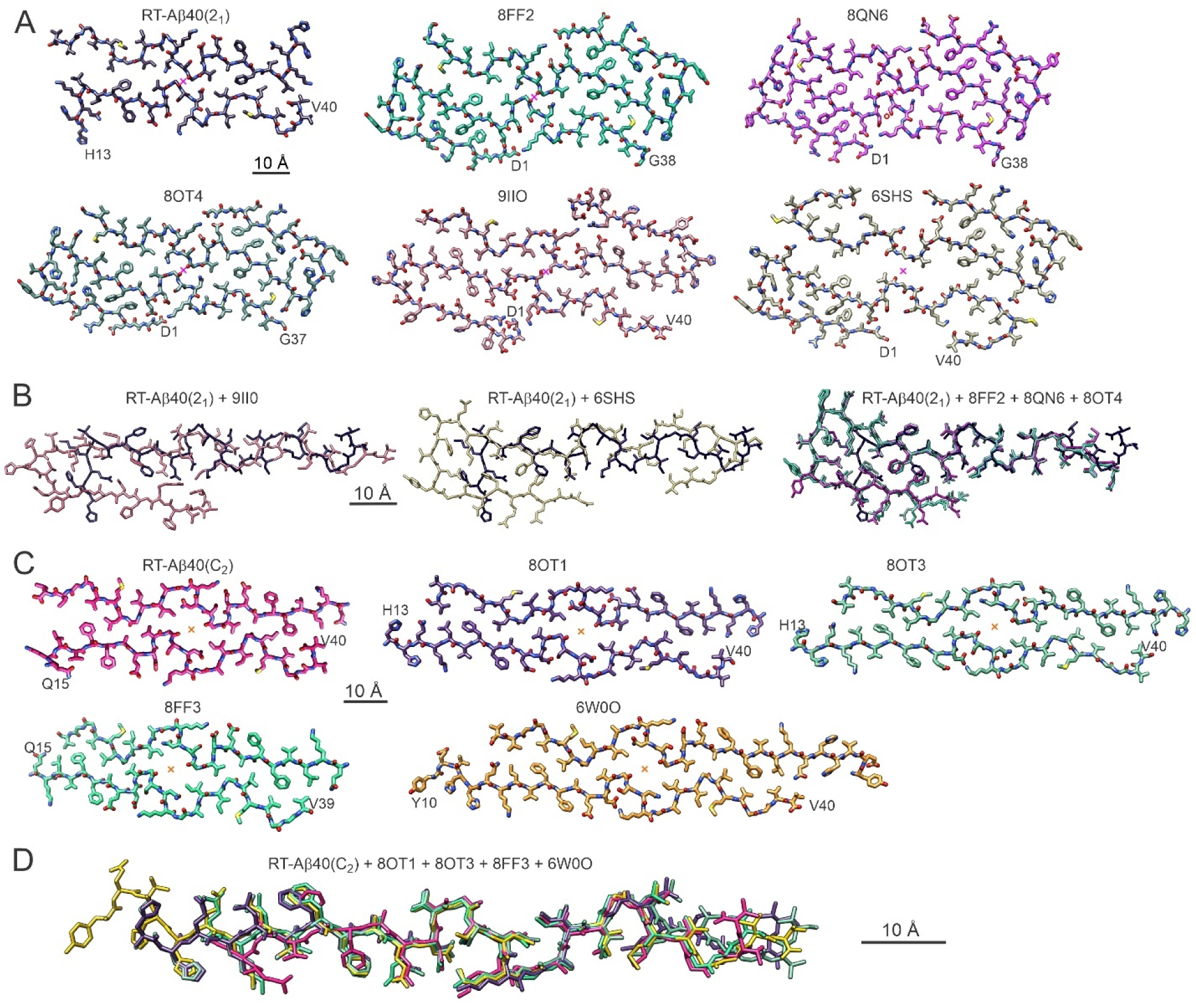
Comparisons of molecular models for RT-Aβ40(2_1_) and RT-Aβ40(C_2_) fibrils with molecular models for previously described Aβ40 fibril polymorphs. (A) Model for RT-A40(2_1_) fibrils and models with the indicated PDB codes. These models share similar structures around their central symmetry axes, indicated by magenta crosses. (B) Alignment of residues 17-36 of one molecule from the RT-Aβ40(2_1_) model (black) with molecules from models with the indicated PDB codes. RMSD values for all non-hydrogen atoms in residues 18-36 are 2.6 Å, 2.7 Å, 2.6 Å, 6.1 Å, and 3.5 Å relative to the 8FF2, 8OT4, 8QN6, 6SHS and 9IIO models, respectively. (C) Model for RT-Aβ40(C_2_) fibrils and models with the indicated PDB codes. These models share similar structures around their central symmetry axes, indicated by orange crosses. (D) Alignment of residues 15-39 of one molecule from the RT-Aβ40(C_2_) model (pink) with molecules from models with PDB codes 8OT1, 8OT3, 8FF3, and 6W0O. RMSD values for all non-hydrogen atoms in residues 15-39 are 3.7 Å, 3.7 Å, 3.0 Å, and 3.18 Å, respectively

**Table 2:** Comparisons of RT-Aβ40(2_1_) and RT-Aβ40(C_2_) fibril structures with structures of previously described Aβ40 fibril polymorphs (represented by their PDB codes).

| Polymorph | Source | Cryo-EM map resolution (Å) | Rise (Å) | Twist (°) | Cross-over distance (nm) | Handedness | Symmetry |
| --- | --- | --- | --- | --- | --- | --- | --- |
| RT-Aβ40(2<sub>1</sub>) | Grown *in vitro* | 2.5 | 2.34 | -181.75 | 24 | right | pseudo-2<sub>1</sub> |
| 8OT4 | Seeded with fibrils extracted from AD meninges | 2.97 | 2.38 | 181.18 | 36 | right | pseudo-2<sub>1</sub> |
| 9IIO | Grown *in vitro* | 3.3 | 2.45 | 179.2° | 55 | left | pseudo-2<sub>1</sub> |
| 8QN6 | AD meninges | 2.4 | 4.87 | -0.79 | 111 | right | C<sub>2</sub> |
| 8QN7 | AD meninges | 2.7 | 2.44 | 179.53 | 93 | right | pseudo-2<sub>1</sub> |
| 8FF2 | Seeded with cerebrovascular amyloid | 2.9 | 2.44 | 181.17 | 38 | right | pseudo-2<sub>1</sub> |
| 6SHS | AD meninges | 4.4 | 2.41 | 181.0 | 43 | right | pseudo-2<sub>1</sub> |
| RT-Aβ40(C<sub>2</sub>) | Grown *in vitro* | 2.7 | 4.69 | -3.55 | 24 | left | C<sub>2</sub> |
| 8OT1 | Grown *in vitro* | 2.59 | 4.763 | -0.782 | 110 | left | C<sub>2</sub> |
| 8OT3 | Grown *in vitro* | 2.73 | 2.372 | 179.526 | 90 | left | pseudo-2<sub>1</sub> |
| 8FF3 | Seeded with cerebrovascular amyloid | 3.1 | 2.44 | 179.75 | 176 | not identified | pseudo-2<sub>1</sub> |
| 6W0O | Seeded with AD cortical homogenate | 2.77 | 2.45 | -180.34 | 130 | left | pseudo-2<sub>1</sub> |

### Comparisons between RT-Aβ40(C_2_) fibrils and previously described polymorphs

As shown in Figs. 4C and 4D, the peptide conformation and inter-subunit contacts in RT-Aβ40(C_2_) fibrils closely resemble conformations and contacts in previously characterized fibril polymorphs with substantially larger cross-over distances (Table 2), including fibrils grown *in vitro* (15) (PDB codes 8OT1 and 8OT3) or grown from tissue-derived seeds (14,20) (PDB codes 8FF3 and 6W0O). To investigate the structural basis for differences in cross-over distances, we focused on comparisons between RT-Aβ40(C_2_) fibrils and the slowly twisting polymorph (PDB code 6W0O) that we have shown previously to be the predominant structure in Aβ40 fibrils prepared by seeded growth from amyloid in cortical tissue of AD patients.

In the slowly-twisting, brain-seeded polymorph, conformational order begins at H14 (20), rather than at L17 as in RT-Aβ40(C_2_) fibrils (Figs. 2H and S7B). Again, the shorter conformationally ordered segment may contribute to the relatively rapid twist in RT-Aβ40(C_2_) fibrils. The two polymorphs also differ in the angle between the major molecular axis direction, defined as the direction from the N-terminal end of the conformationally ordered segment to the

C-terminal end, and the fibril growth direction. As shown in Fig. 5A, this difference produces a larger angle between major molecular axes in the two cross-β subunits, i.e., a more pronounced “herringbone” pattern, in the RT-Aβ40(C_2_) fibrils. The relatively small angle in the PDB 6W0O polymorph (approximately 2°) favors its observed pseudo-2_1_ symmetry, which allows interdigitation of sidechains from the two subunits at the inter-subunit interface. The larger angle in the RT-Aβ40(C_2_) fibrils (approximately 7°) allows similar interdigitation of sidechains from the two subunits in C_2_ symmetry.

**Figure 5:**
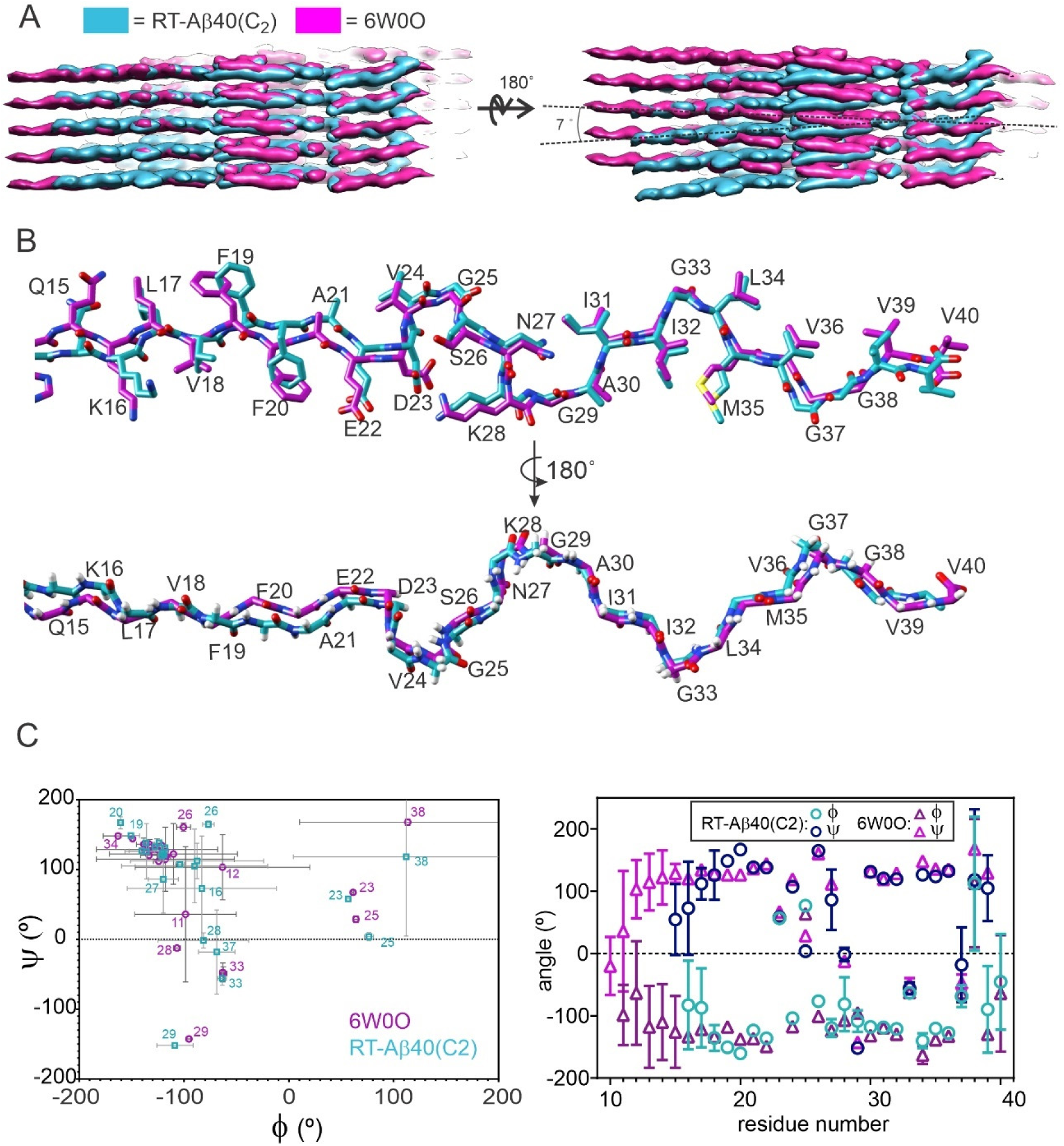
Comparisons of cryo-EM densities and molecular models for rapidly twisting RT-Aβ40(C_2_) fibrils and slowly twisting brain-seeded fibrils (PDB code 6W0O). (A) Side views of the overlaid density maps. Optimal alignment of densities in one cross-β subunit (left) results in an approximate 7° misalignment of densities in the other subunit (right). (B) Superposition of the two molecular models, showing differences in backbone conformations (lower superposition) and sidechain conformations (upper superposition), especially in residues 15-25. (C) Comparisons of backbone φ and ψ torsion angles, represented either as a Ramachandran plot (left) or as a plot of angles vs. residue number (right). Angle values are averages over the six molecules and 10 molecular models in the structure bundles, with standard deviations indicated by error bars.

Despite the factor of 5.4 difference in twist period (Table 2), conformations of residues 17-39 are very similar in the two polymorphs, as shown in Fig. 5B. Conformational differences in molecular models are largest in residues 18-21, including differences in sidechain conformations of V18, F19, and F20 as well as displacements between backbone atoms. Plots of backbone φ and ψ torsion angles in Fig. 5C, averaged over molecules in the bundles of structures generated by Xplor-NIH calculations, show that these angles are similar but not identical. Thus, it appears that the more rapid twist of RT-Aβ40(C_2_) fibrils arises from a combination of a shorter conformationally ordered segment, a larger deviation from perpendicularity between major axis directions and the fibril growth direction, and subtle differences in conformation that are spread over multiple sites. The more rapid twist is not attributable to large conformational differences at a specific site or to the presence or absence of a specific intermolecular interaction.

To gain more insight into the properties of these rapidly twisting and slowly twisting polymorphs, we performed 200 ns molecular dynamics (MD) simulations in explicit H_2_O solvent with 100 mM NaCl at 310 K. Analyses of the MD trajectories showed that M35 sidechains were dynamically disordered in both polymorphs, with large conformational variations on the 1 ns time scale (Fig. S11A). In RT-Aβ40(C_2_) fibrils, F19 and F20 sidechains were relatively immobile, but adopted more than one conformation and switched between conformations on 10-100 ns time scales (Fig. S11B). In the slowly twisting polymorph, F19 sidechains were immobile and conformationally ordered, while F20 sidechains were highly dynamic (Fig. S11C). Differences in F19 and F20 sidechain ordering and dynamics may correlate with the conformational differences discussed above. D23-K28 salt bridge interactions were stable in both polymorphs, but with a longer average sidechain-sidechain distance in RT-Aβ40(C_2_) fibrils (3.49 Å) than in the slowly twisting fibrils (3.32 Å) (Fig. S11C).

Water molecules occupied central channels in MD simulations on both RT-Aβ40(C_2_) fibrils and slowly twisting, brain-seeded fibrils. Within these channels, water molecules were localized near the peptide backbone at S26 and the sidechain of I31 (Fig. S12A). Aside from the central channels, water molecules were largely excluded from both fibril cores. The MD simulations also showed that Na^+^ ions tended to associate with negatively charged sidechains of E22 in RT-Aβ40(C_2_) fibrils, while Cl^-^ ions tended to associate with the positively charged amino groups of K28 sidechains. In the slowly twisting fibrils, Na^+^ ions associated with sidechains of both E22 and D23, association of Cl^-^ ions with K28 sidechains was less pronounced, and Cl^-^ ions also associated with sidechains of K16, which were ordered in the slowly twisting fibrils but disordered in RT-Aβ40(C_2_) fibrils (Fig. S12B). Differences in ion distributions may reflect differences in electrostatic interactions that contribute to differences in fibril twist.

## Discussion

Results described above show that three structurally distinct, rapidly twisting Aβ40 fibril polymorphs with nearly identical cross-over distances can grow simultaneously *in vitro*. Two of these, the RT-Aβ40(2_1_) and RT-Aβ40(C_2_) polymorphs, have core structures that resemble those of previously reported Aβ40 fibril polymorphs with significantly longer cross-over distances (Fig. 4 and Table 2). The third polymorph, RT-Aβ40(C_1_), is unique in that the two cross-β subunits in its core are not related by symmetry. Such an asymmetric structure has not been reported previously for wild-type Aβ40 or Aβ42 fibrils.

A structure for E22G-Aβ40 fibrils extracted from cortical tissue of a familial AD patient, described by Yang *et al.* (18) (PDB code 8BG0), contains peptide molecules with two different conformations. However, these molecules occur as two inequivalent pairs of subunits, arranged to produce overall C_2_ symmetry in the E22G-Aβ40 fibril structure.

The RT-Aβ40(2_1_) polymorph belongs to a class of polymorphs defined by inter-subunit G25-S26 and V24-I32 contacts and solvent exposure of I31 sidechains (Fig. 2E). This class includes structures with PDB codes 8FF2, 8QN6, and 8OT4 and may also include PDB codes 9IIO and 6SHS (Fig. 4A). The RT-Aβ40(C_2_) polymorph belongs to a class of polymorphs defined by inter-subunit V24-I31 and V24-G33 contacts and solvent exposure of I32 sidechains (Fig. 2I). This class includes structures with PDB codes 8OT1, 8OT3, 8FF3, and 6W0O (Fig. 4C) as well as PDB code 8OVK. The RT-Aβ40(C_1_) polymorph can be placed in the first of these two classes (Fig. 3E), although the absence of two-fold symmetry, major conformational differences in residues 33-38 of the upper subunit (pink density in Fig. 3), and many additional conformational differences distinguish RT-Aβ40(C_1_) fibrils from RT-Aβ40(2_1_) fibrils.

The more rapid twist of RT-Aβ40(2_1_) and RT-Aβ40(C_2_) fibrils, compared with structurally similar polymorphs, can not be attributed to conformational differences at specific sites in the structurally ordered cores. Instead, more rapid twist appears to correlate with shorter structurally ordered segments. Simple geometric considerations may contribute to this correlation, based on the fact that intermolecular distances are greater at the outer edge of a twisting, cross-β amyloid structure than at the center. Given that a cross-β structure requires a 0.48 ± 0.01 nm distance between peptide molecules in the fibril growth direction, a cross-over distance equal to *d* requires an angle *ρ* = 0.48*π* / *d* between successive molecules in the cross-β structure. If the conformationally ordered segments of peptide molecules have extended conformations with lengths equal to *L*, then the intermolecular distances at the N- and C-terminal ends of the ordered segments become 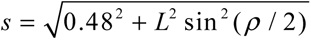. Thus, a length of 7.3 nm (residues 14-40 in PDB code 6W0O) and a cross-over distance of 25 nm creates intermolecular distances equal to 0.528 nm at the N- and C-terminal ends, while a length of 5.3 nm (residues 17-37 in RT-Aβ40(C_2_) fibrils) and a cross-over distance of 25 nm creates intermolecular distances equal to 0.506 nm. Large intermolecular distances that would be created by large molecular lengths and small cross-over distances may prevent the formation of stable intermolecular hydrogen bonds, thereby limiting the fibril twist rate. A related argument was given by Meinhardt *et al.* in an early cryo-EM study of Aβ40 fibrils (4).

An important issue in the field of amyloid research is whether fibril polymorphs that are purified directly from biological tissue depend on chemical cofactors, interfaces, or other aspects of the biological environment that are absent when fibrils are grown *in vitro*, thus leading to fundamental differences between fibrils in tissue and fibrils generated in the laboratory. A related issue is whether the molecular structures of polymorphs that develop in tissue can be propagated or amplified by seeded growth, *i.e.*, by addition of monomeric peptides or proteins to solutions that contain short fragments of fibrils from tissue(3,23,36). Aβ40 fibril structures with PDB codes 6SHS and 8QN6 come from cryo-EM measurements on fibrils that were extracted from meninges of AD patients (10,12). Although the RT-Aβ40(2_1_) polymorph lacks conformational order in residues 1-12 and has a different conformation in residues 33-40, the conformation of residues 18-32 in RT-Aβ40(2_1_) fibrils is nearly identical to that in the Aβ40 fibrils from meninges (Fig. 4B). Thus, while molecular structures of Aβ40 fibrils grown *in vitro* in simple aqueous buffers may not be the same as structures of fibrils that develop in tissue, substantial aspects of these structures can be quite similar. Moreover, the sense of twist is right-handed in RT-Aβ40(2_1_) fibrils, as it is in fibrils extracted from meninges, showing that right-handed twist is not a unique characteristic of fibrils that develop in tissue. Experiments by Pfeiffer *et al*. (15) and Fu *et al.* (*14*) have also shown that the structure of Aβ40 fibrils from meninges can be propagated faithfully by seeded fibril growth, leading to the structures with PDB codes 8FF2 and 8OT4 (Fig. 4A).

As discussed in detail above, RT-Aβ40(C_2_) fibrils closely resemble fibrils with PDB code 6W0O, which were prepared by seeded growth from amyloid in cortical tissue homogenate from AD patients (20). RT-Aβ40(C_2_) fibrils also closely resemble fibrils with PDB code 8FF3, which were produced by seeded growth from microdissected cerebral amyloid angiopathy tissue (14). To our knowledge, fibril polymorphs that are structurally similar to RT-Aβ40(C_2_) fibrils (or to fibrils with PDB codes 6W0O or 8FF3) have not yet been found in cryo-EM studies of fibrils that were directly extracted from AD brain tissue. This may mean that such polymorphs do not exist in AD brain tissue. However, it is also possible that such polymorphs do exist in AD brain tissue but are difficult to isolate with sufficient purity for high-resolution structural characterization cryo-EM.

Finally, S-shaped peptide conformations have been found in Aβ42 fibrils grown *in vitro* (8,25–27) and in Aβ42 fibrils that were isolated from AD brain tissue (11). S-shaped peptide conformations in cryo-EM-based structures of wild-type Aβ40 fibrils have not been reported and were not found in the present studies of rapidly twisting Aβ40 fibrils. On the other hand, an ssNMR-based model for fibrils formed by E22Δ-Aβ40 does include S-shaped peptide conformations similar to those in Aβ42 fibrils (24), as do models for E22G-Aβ40 fibrils extracted from transgenic mouse and familial AD brain tissue (18,19). A ssNMR-based model for brain-seeded fibrils from an atypical AD patient includes related peptide conformations (23). Thus, future studies may further expand the range of known Aβ40 and Aβ42 fibril structures.

## Materials and Methods

### Aβ40 fibril preparation

Aβ40, synthesized and purified as previously described (37), was dissolved in dimethyl sulfoxide (DMSO) to create a stock solution with 6.7 mM Aβ40 concentration. To find a condition that produces a large population of rapidly twisting fibrils, fibrils were grown quiescently under various conditions of temperature, pH, peptide concentration, buffer and salt concentration, and incubation time. Briefly, aliquots of the DMSO stock solution were diluted to the final desired Aβ40 concentration (50-200 μM) into the desired buffer at room temperature or 37° C and vortexed immediately to prevent the peptide aggregation. Solutions were then incubated without shaking or rocking for a minimum of 6 days. Each sample was then imaged using negative stain. Figure S1 shows TEM images of fibrils that were grown under some of the tested conditions. Fibrils that were grown from 200 μM Aβ40 peptide concentration in 10 mM sodium phosphate buffer, pH 7.4, 0.01% sodium azide, at 37° C showed the highest population of rapidly twisting fibrils. More TEM images of this sample are shown in Figure S2.

### Negative stain transmission electron microscopy

For negative stain TEM image, Quantifoil R2/1 grids with 2 nm carbon films and lacey carbon grids from Electron Microscopy Sciences with hand-made carbon films were used. Samples were gently vortexed and diluted in H_2_O to make a final monomer concentration around 25 μM. 10 µl aliquots of diluted samples were deposited on glow discharged grids. After 2 min adsorption, grids were blotted, rinsed with 10 µl H_2_O, blotted, and stained with 10 µl of 2% w/v uranyl acetate. Uranyl acetate was blotted after 1-2 minutes of staining, and grids were dried in air. Images were acquired with an FEI Morgagni electron microscope at 80 keV with a side-mounted Advantage HR camera (Advanced Microscopy Techniques).

### Cryo-electron microscopy grid preparation and data acquisition

Grids (Quantifoil Cu400 R1.2/1.3) were glow-discharged for 45 s in a PELCO easiGlow instrument. Aβ40 fibril solutions with high populations of rapidly twisting fibrils were diluted by factors of 2 or 4 in H_2_O. 3 µl of the diluted sample was deposited on grids in 100% humidity, then blotted for 2.5 s or 3 s with force level 1 in a Vitrobot IV plunge freezer (ThermoFisher Scientific).

Grids were vitrified by plunging into liquid ethane. Grids prepared with 4X dilution were used for acquiring the images. Those that were prepared with 2X dilution were found to be overcrowded.

Grids were then screened using a ThermoFisher Glacios microscope that operates at 200 keV and is equipped with a Gatan K3 camera. Final images were acquired on a ThermoFisher Titan Krios G4 microscope operating at 300 keV. Parameters used for acquiring the images are listed in Table 1. Images were recorded automatically using Serial EM and EPU software.

### Cryo-EM image processing and helical reconstruction

All early processing steps were done in RELION 5.0 (29). Final 3D reconstruction steps were done with a modified version of RELION 4.0 (20). Micrographs acquired with SerialEM and EPU software were combined before any processing. Motion correction on all 6791 micrographs was done in RELION. After contrast transfer function (CTF) estimation, manual particle picking was performed on a small set of rapidly twisting fibrils. The box size of 1000 pixels was rescaled to 500 pixels (0.82 Å pixel size), with an inter-box distance of 48 Å. After two rounds of 2D classification and selection of well-resolved 2D class average images, 12972 particles from 146 micrographs were used to perform Topaz training (38). Particles were then picked automatically using the trained Topaz model. A total of 3022787 particles were extracted.

After one round of 2D classification, class averages containing distinguishable particles were selected (1217380 particles). After another round of 2D classification, 2D class average images showing rapidly twisting morphologies were selected, and 2D classes showing similar morphologies were assigned to RT-Aβ40(2_1_), RT-Aβ40(C_2_), and RT-Aβ40(C_1_) polymorphs. Well-resolved 2D class averages were selected and their psi angles (in-plane orientations) were adjusted, using a MATLAB script as previously described to align the fibril growth axes with the x axis of the 2D class (20).

Before creating an initial 3D model, the 2D class averages were used to manually estimate the rotation angles, assuming a 180° rotation about the fibril growth direction between the narrowest points of the fibrils in the 2D class average images. The estimated rotation angles were assigned to the RELION AngleRot variable for particles in each 2D class as previously described (20). The X,Y coordinate values were modified after 2D classification to place the particles back in their original position along the fibril growth axis. These particles were then used to generate an initial 3D model by running one iteration of 3D classification with a single class and with a reference model generated by the “3D initial reference” function of RELION 5.0. Initial helical twist and rise values were estimated from the 2D class averages and were later refined during 3D refinement using local helical symmetry searches.

3D classification was performed separately for sets of particles that were assigned to RT-Aβ40(2_1_), RT-Aβ40(C_2_), and RT-Aβ40(C_1_) polymorphs. Four 3D classes were allowed for each polymorph. For RT-Aβ40(2_1_) and RT-Aβ40(C_2_) fibrils, density maps for all 3D classes were similar after 3D classification. For RT-Aβ40(C_1_) fibrils, one of the 3D classes appeared to be different from the other three after 3D classification. The distinct 3D class became the RT-Aβ40(C_1_)′ polymorph. After several rounds of 3D refinement, 2D classification, and selection of particles from the well resolved classes, particles from filaments shorter than 5 particles were removed. Before the final refinement, rotation and psi angles were modified using MATLAB script described before. CTF refinement and Bayesian polishing were then performed and final 3D refinements were run with and without additional symmetry.

In 2D classification of the RT-Aβ40(C_1_) particles, intensity modulation suggestive of a repeat distance approximately equal to 9.4 Å was observed. Therefore, 3D refinement was performed with this repeat distance. In the final 3D density map, only one of the weak extra density layers (below the lower subunit) has a clear 9.4 Å periodicity, rather than 4.7 Å. The core structure has 4.7 Å periodicity.

### Model building and Xplor-NIH calculations

Attempts to fit the Aβ40 peptide sequence into 3D density maps for rapidly twisting Aβ40 fibrils using the ModelAngelo feature of RELION 5.0 (39) did not generate a continuous model. Initial models for one Aβ40 in the 3D density maps were therefore constructed with Coot software (31). The residues included in molecular models for each polymorph are given in Table 1. Six copies of these manually constructed models were inserted into 3D density maps with specific symmetry and handedness with the “Fit in Map” function of UCSF Chimera (40), creating models for fibril segments with two cross-β subunits and three repeats along the fibril growth direction. Further refinement and energy minimization were done in Xplor-NIH (32) with the parameters specified in Table S1. Briefly, all non-hydrogen atoms with clearly resolved densities were restrained within the cryo-EM density map with the probDistPot energy function of Xplor-NIH, and molecular conformations within each cross-β subunit were kept nearly identical to each other with the NCS and DistSymmPot energy functions. Backbone heavy atoms of each chain were restrained to within 2.5 Å of their original position with the PosDiffPot function. Annealing in Xplor-NIH was performed from 3000 K to 25 K with decrements of 12.5 K and 5000 simulation steps at each temperature. After annealing, final energy minimizations were performed in torsion angle and Cartesian coordinates.

Multiple rounds of Xplor-NIH calculations were performed for each polymorph (6 for RT-Aβ40(2_1_) fibrils, 3 for RT-Aβ40(C_2_) fibrils, and 5 for RT-Aβ40(C_1_) fibrils, with 40 independent annealing simulations in each run). The lowest-energy structure from each round was used as the initial structure for the next round until the final round, which generated a set of structures without significant steric clashes or restraint violations. For each polymorph, 10 structures with no violations according to Xplor-NIH and no significant clashes according to the PDB validation website and Molprobity were selected and deposited in the Protein Data Bank. These structure bundles are shown in Fig. S5.

### All-atom molecular dynamics simulations

To compare the molecular interactions and dynamics of RT-Aβ40(C_2_) fibrils and the brain-derived, slowly twisting Aβ40 fibril (PDB code 6W0O), we performed molecular dynamic simulations using NAMD software (41). Using the final Xplor-NIH-calculated models for each polymorph, we constructed fibril segments containing 14 peptide molecules in UCSF Chimera software, with two symmetry-related cross-β subunits and seven repeats in each subunit. N-terminal segments that were not included in the Xplor-NIH calculations were added to the fibrils with initial β-sheet structures. To randomize the conformations of the N-terminal segments, preliminary MD simulations were first run in vacuum at 600 K for 50 ns, then 310 K for 50 ns. Residues 11-40 and 16-40 of the brain-derived fibril segment and the RT-Aβ40(C_2_) fibril segment, respectively, were fixed in place during these preliminary simulations and the dielectric constant was set to 100.

Final structures from the preliminary simulations were then solvated and neutralized in 100 mM NaCl in a 62.7 Å × 133.7 Å × 59.45 Å (RT-Aβ40(C_2_) fibril) or 55.15 Å × 153.57 Å × 51.90 Å (brain-derived fibril) periodic box in VMD software (42). To prevent fraying of the ends of the fibril segments during MD simulations, positions of the C_α_ atoms of residues 17-40 in the RT-Aβ40(C_2_) fibril or residues 11-40 in the brain-derived fibril were constrained using a harmonic constraint scaling factor of 0.05 kcal/mol-Å^2^. Simulations were performed with CHARMM36 force field parameters (43), Langevin dynamics at 310 K, constant pressure, and periodic boundary conditions. Simulations for both fibrils were run for 200 ns, with coordinates being saved every 0.2 ns (1000 movie frames).

Plots of torsion angles and interatomic distances in Fig. S11 were prepared in GraphPad Prism, using values that were extracted in VMD for the three central repeats. To prepare Fig. S12, ordered residues were aligned to the last frame. The display was clipped in VMD to contain only the three central repeats. Images from 375 frames, showing protein atoms, sodium ions, chloride ions and water oxygen atoms individually were summed in FIJI software and superimposed.

## Data and software availability

Scripts used in cryo-EM density map calculations, scripts used in Xplor-NIH calculations, scripts and files used in MD simulations, trajectories from MD simulations, density maps with disfavored twist handedness, and the corresponding molecular models are available at https://doi.org/10.17632/j7k4ksx5fc.1.

## Acknowledgements

This research was supported by the Intramural Research Program of the National Institute of Diabetes and Digestive and Kidney Diseases (NIDDK) within the National Institutes of Health (NIH). The contributions of the NIH authors are considered Works of the United States Government. The findings and conclusions presented in this paper are those of the authors and do not necessarily reflect the views of the NIH or the U.S. Department of Health and Human Services. We thank Dr. Wai-Ming Yau for synthesis and purification of Aβ40 peptides. This work utilized the computational resources of the NIH HPC Biowulf cluster (https://hpc.nih.gov), as well as instrumentation of the NIDDK Cryo-Electron Microscopy Core and the NIH Multi-Institute Cryo-Electron Microscopy Facility (MICEF).

## Author contributions

MGL and RT designed the research; MGL performed experiments; MGL and KRT analyzed data; MGL and RT wrote the paper. All authors gave approval to the final version of the paper.

## Supplemental information

Movies 1-6 and Supporting Information (containing Table S1, movie captions, and Figs. S1-S12).

## Notes

https://doi.org/10.17632/j7k4ksx5fc.1

### Competing Interest Statement

The authors have declared no competing interest.

