## Supporting figures and tables for "Polymorphic structures of rapidly twisting 40-residue amyloid-β fibrils"

Motahareh G. Larimi, Kent R. Thurber, and Robert Tycko
Laboratory of Chemical Physics
National Institute of Diabetes and Digestive and Kidney Diseases
National Institutes of Health
Bethesda, MD 20892-0520

**Movie captions**

Movie 1: 200 ns MD simulation of a RT-Aβ40($C_2$) fibril segment in explicit water with 100 mM NaCl at 310 K. A central slab containing 6 peptide molecules is shown, viewed along the fibril growth direction. $Na^+$ and $Cl^-$ ions are represented by magenta and yellow spheres, respectively.

Movie 2: Expanded view from the MD simulation of a RT-Aβ40($C_2$) fibril segment, showing the dynamics of F19 and F20 sidechains from one subunit (cyan and pink, respectively) and M35 sidechains from the other subunit (yellow).

Movie 3: Same as Movie 2, but with the opposite choice of subunits.

Movie 4: 200 ns MD simulation of a PDB 6W0O fibril segment in explicit water with 100 mM NaCl at 310 K. A central slab containing 6 peptide molecules is shown, viewed along the fibril growth direction. $Na^+$ and $Cl^-$ ions are represented by magenta and yellow spheres, respectively.

Movie 5: Expanded view from the MD simulation of a PDB 6W0O fibril segment, showing the dynamics of F19 and F20 sidechains from one subunit (cyan and pink, respectively) and M35 sidechains from the other subunit.

Movie 6: Same as Movie 5, but with the opposite choice of subunits.

**Table S1:** Structural restraints in Xplor-NIH calculations

| Restraint potential | Scale factor | | |
|---|---|---|---|
| | RT-Aβ40($2_1$) | RT-Aβ40($C_2$) | RT-Aβ40($C_1$) |
| probDistPot | 10 | 10 | 10 |
| NCS | 100 | 100 | 100 |
| DistSymmpot | 100 | 100 | 100 |
| Repelpot[a] | 0.004 to 4 | 0.004 to 4 | 0.004 to 4 |
| TorsionDB[a] | 0.002 to 2 | 0.002 to 2 | 0.002 to 2 |
| BOND | default | default | default |
| ANGL[a] | 0.4 to 1 | 0.4 to 1 | 0.4 to 1 |
| IMPR[a] | 0.1 to 1 | 0.1 to 1 | 0.1 to 1 |
| Residues included in probDistPot | 14-39 | 16-40[b] | 14-37 |
| Number of independent calculations in final round | 40 | 40 | 40 |

[a]Scale factor incremented from minimum to maximum value during annealing.
[b]Sidechains of residues 16, 17, and 40 not included in probDistPot term.

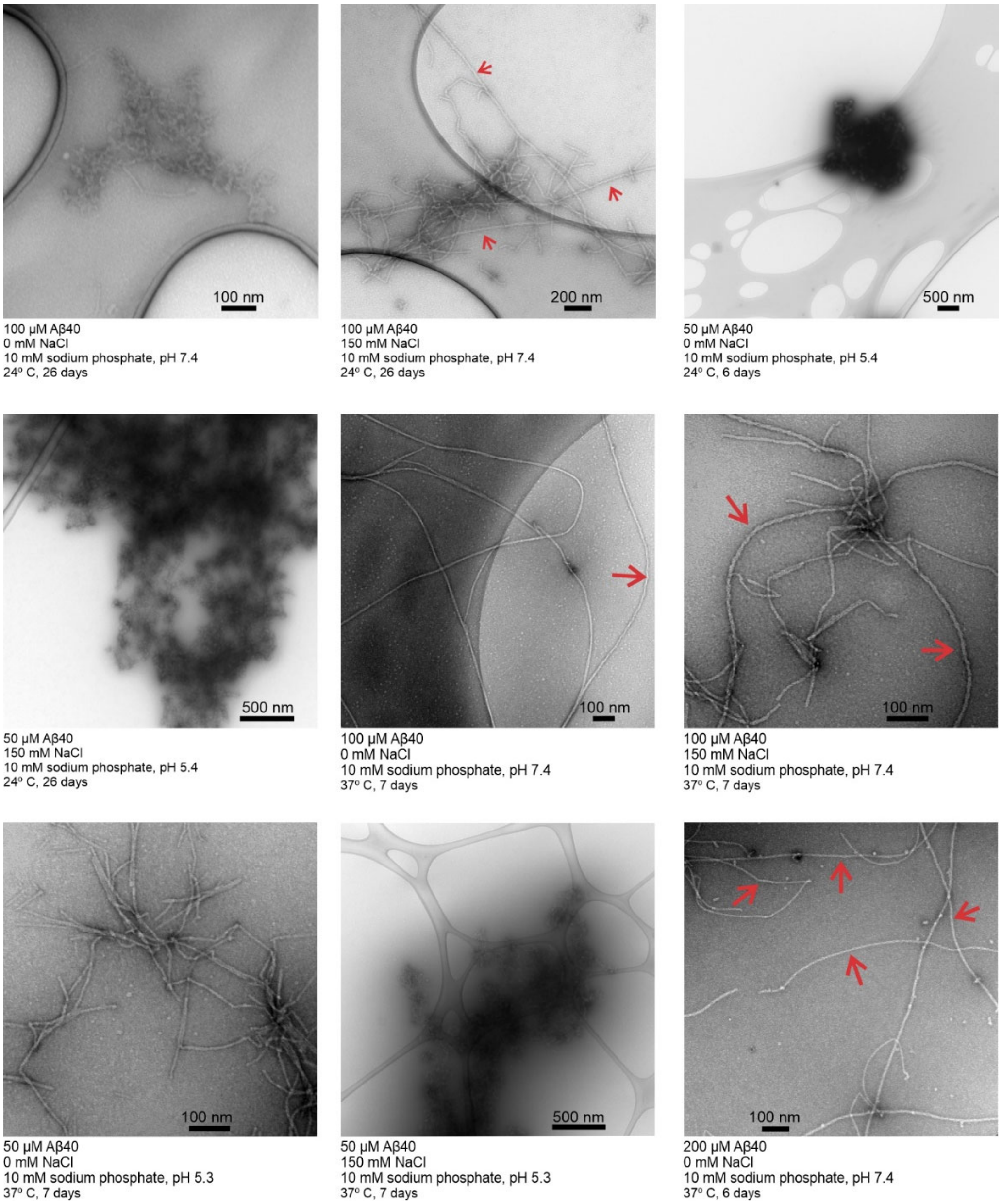


Figure S1: Negative-stain TEM images of Aβ40 fibrils grown in vitro under various conditions. Images were recorded after quiescent incubation for the indicated time periods at the indicated peptide concentrations, NaCl and buffer concentrations, pH values, and temperature. Red arrows indicate rapidly twisting polymorphs.

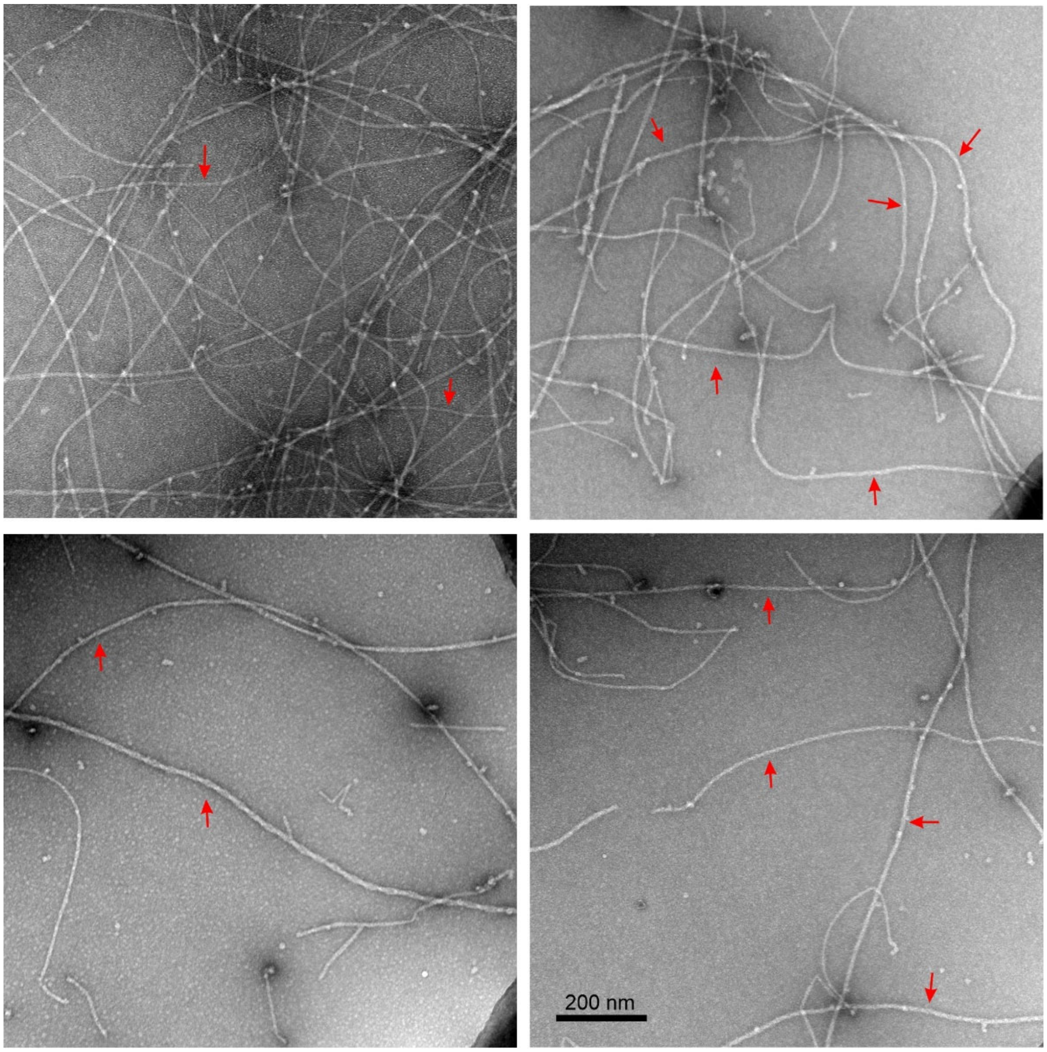


Figure S2**:** Negative-stain TEM images of the Aβ40 fibril sample used for cryo-EM measurements. Fibrils were grown quiescently at 37° C, with 200 μM Aβ40, 10 mM sodium phosphate buffer, pH 7.4, without addition of NaCl. Red arrows indicate rapidly twisting polymorphs.

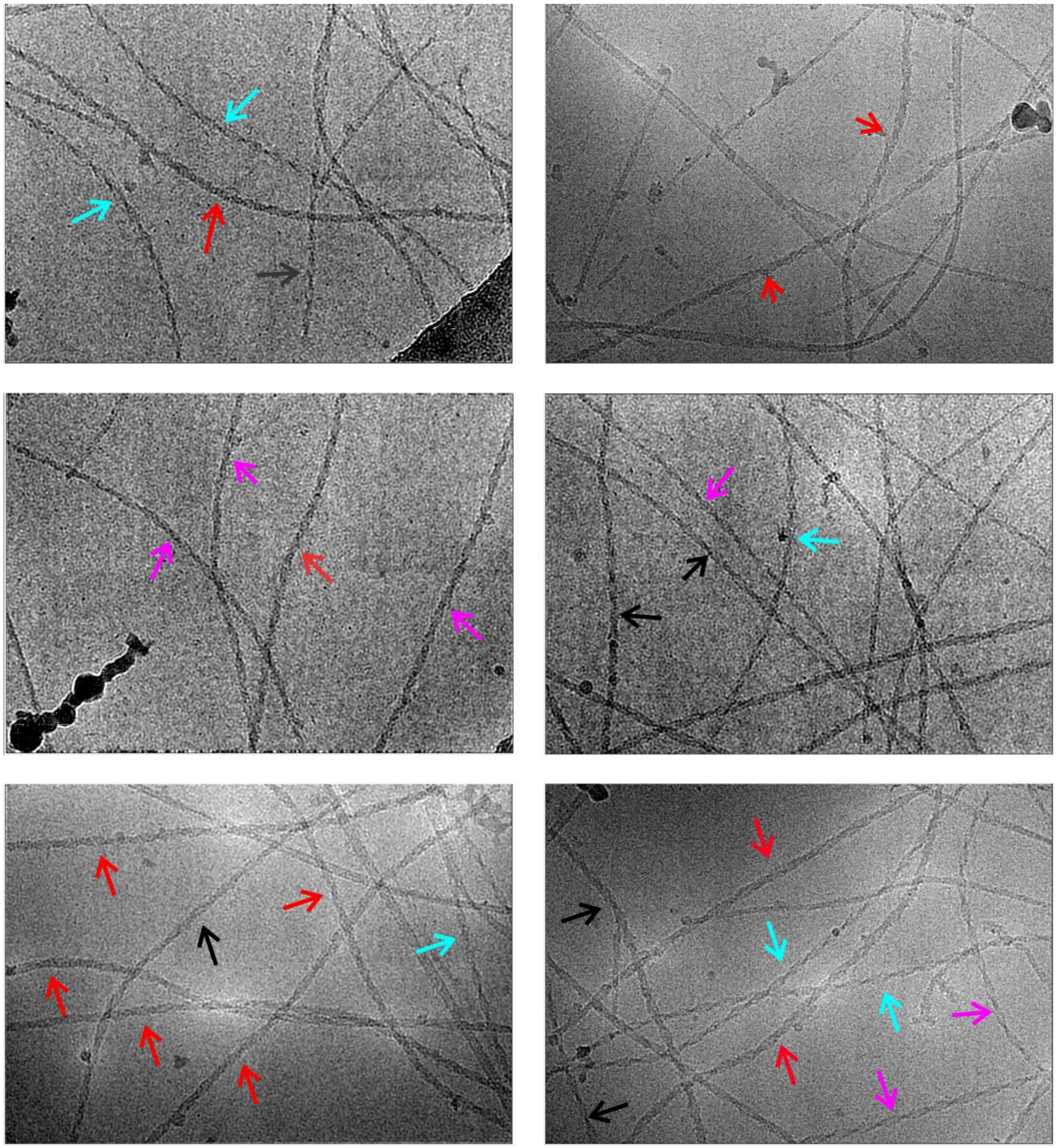

Figure S3: Additional cryo-EM images of fibrils that were grown quiescently at 37° C, with 200 μM Aβ40, 10 mM sodium phosphate buffer, pH 7.4, without addition of NaCl. Cyan, red, and magenta arrows indicate RT-Aβ40($2_1$), RT-Aβ40($C_2$), and RT-Aβ40($C_1$) fibrils, respectively. Black arrows indicate rapidly twisting fibrils that could not be assigned unambiguously to specific polymorphs.

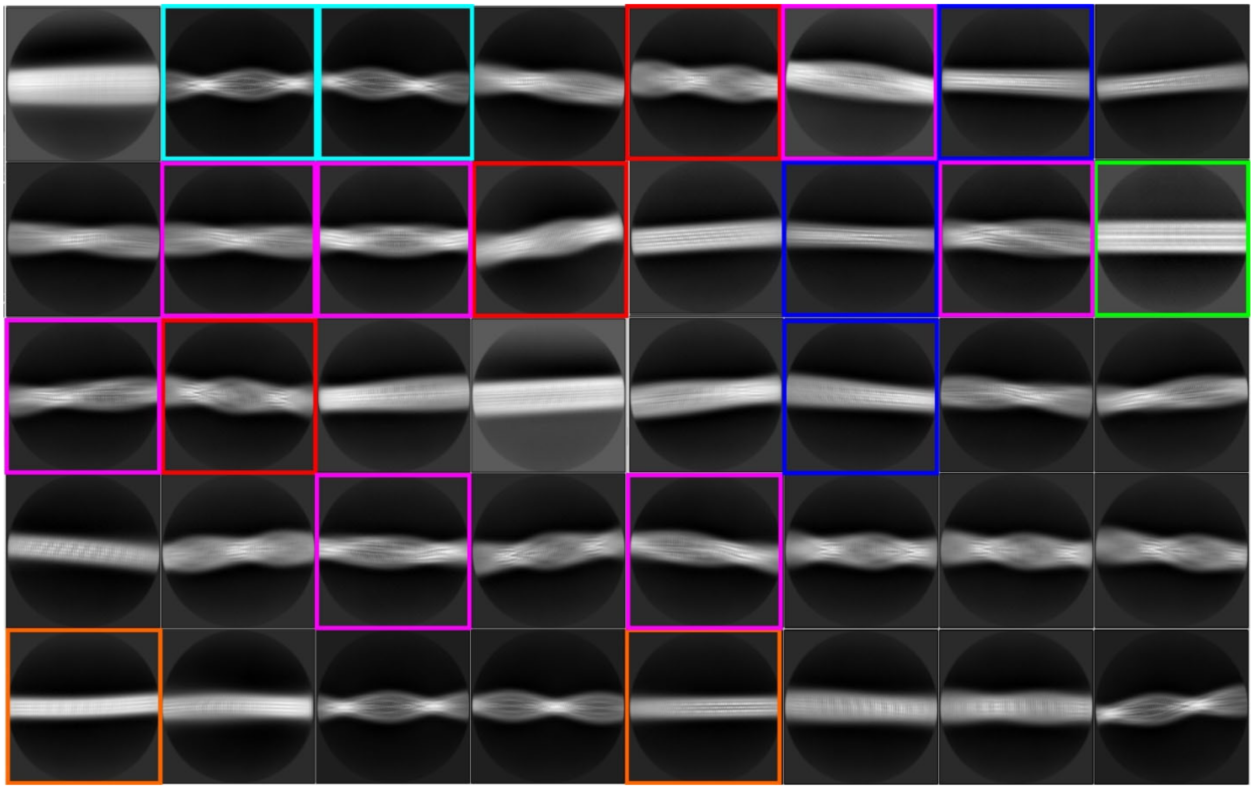

Figures S4: 2D class average images from cryo-EM images of Aβ40 fibrils. The 2D classes show the presence of six polymorphs in this sample. Examples of 2D classes for RT-Aβ40($2_1$), RT-Aβ40($C_2$), and RT-Aβ40($C_1$) fibrils are enclosed in cyan, red and magenta boxes. 2D classes for two polymorphs with no visible twist are enclosed in orange and green boxes. 2D classes for a previously described slowly twisting fibril (PDB code 6W0O) is enclosed in blue boxes. Box sizes are 410 Å.

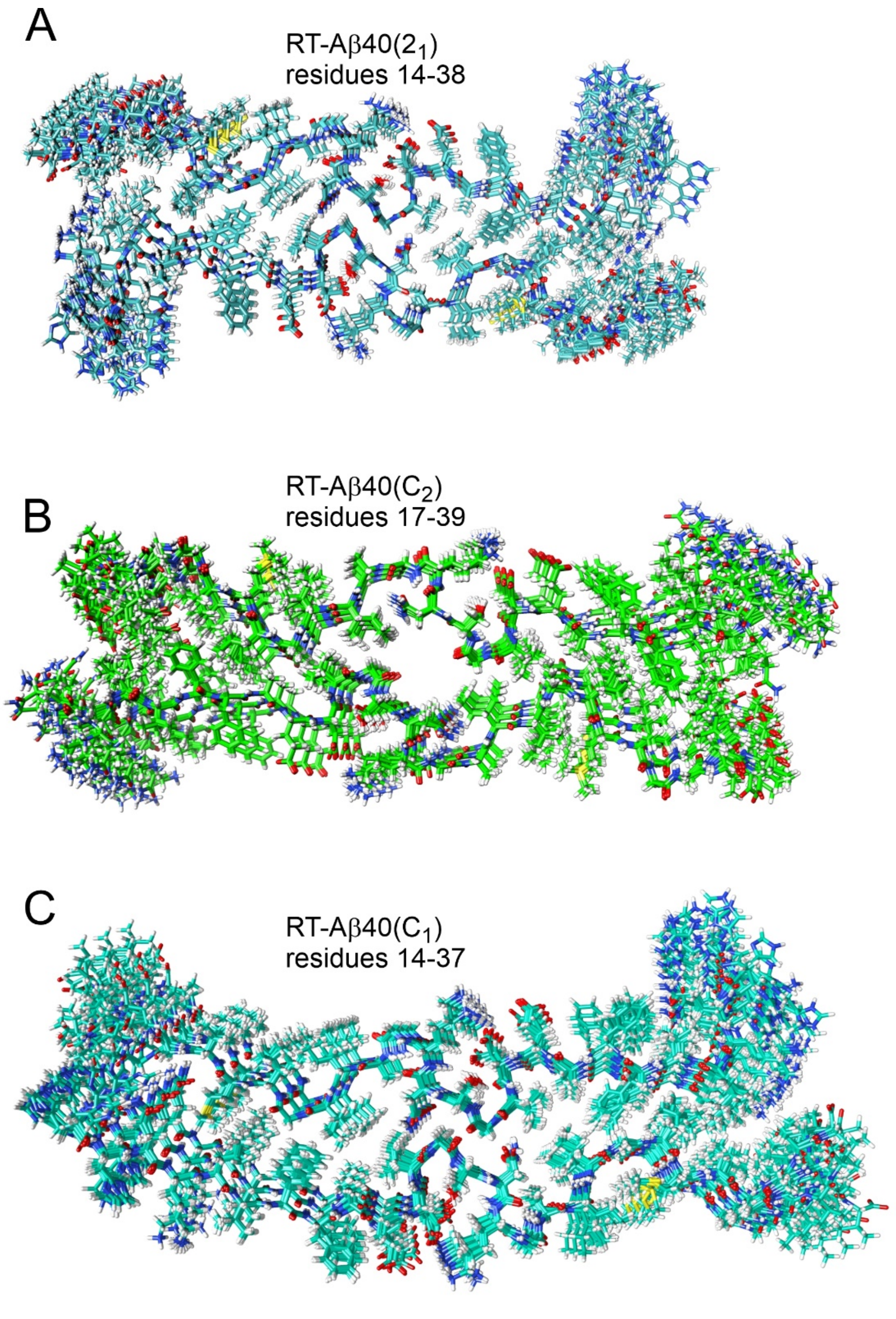


Figure S5: Bundles of 10 independent structural models calculated by Xplor-NIH for (A) RT-Aβ40($2_1$), (B), RT-Aβ40($C_2$), and (C) RT-Aβ40($C_1$) fibrils.

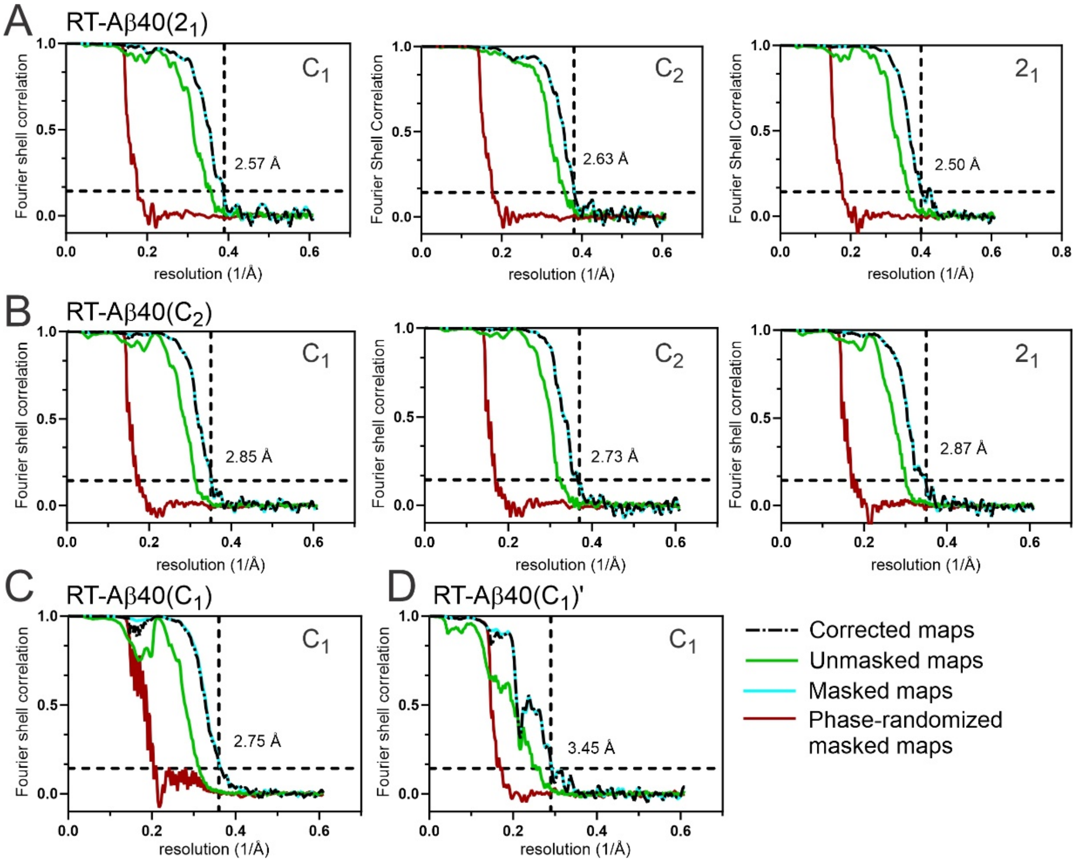


Figure S6: Fourier shell correlation (FSC) curves of 3D density maps calculated for four rapidly twisting Aβ40 fibril polymorphs. FSC curves are shown for unmasked (solid green lines), masked (solid cyan lines), phase-randomized (solid red lines), and corrected (black dash-dot lines) maps. (A) Results for RT-Aβ40($2_1$) fibrils from calculations with $C_1$ symmetry and calculations with $C_2$ symmetry or quasi-$2_1$ symmetry imposed. (B) Results for RT-Aβ40($C_2$) fibrils from calculations with $C_1$ symmetry and with $C_2$ symmetry or quasi-$2_1$ symmetry imposed. (C,D) Results for RT-Aβ40($C_1$) and RT-Aβ40($C_1$)′ fibrils with $C_1$ symmetry.

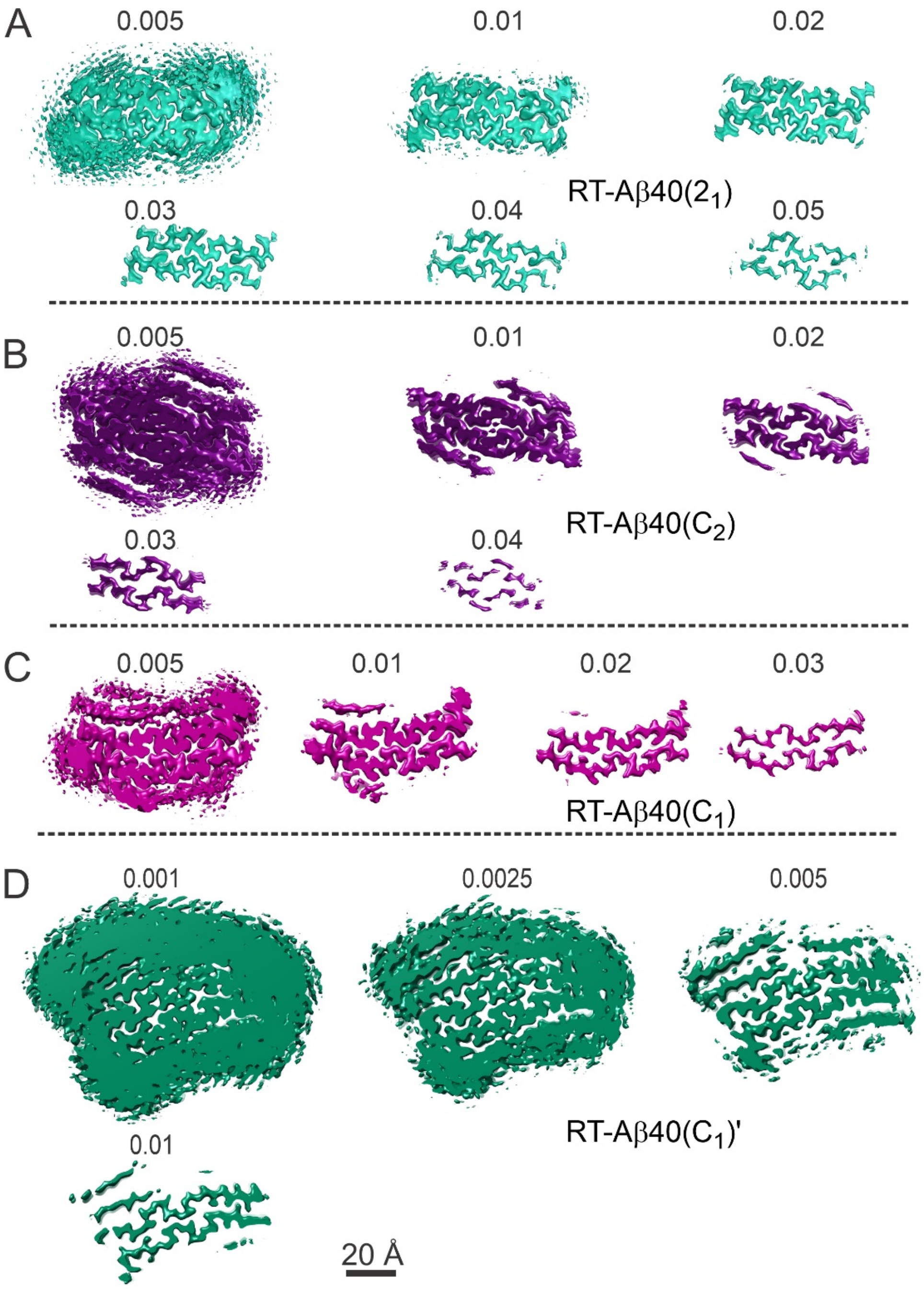


Figure S7: Cross-sectional views of density map surfaces at different contour levels. (A-D) Surfaces of density maps for RT-Aβ40($2_1$), RT-Aβ40($C_2$), RT-Aβ40($C_1$), and RT-Aβ40($C_1$)′ fibrils at the indicated threshold levels, as reported by Chimera software. Plots in panels B, C, D (but not panel A) show additional layers of density above and below the two central cross-β subunits. For RT-Aβ40($C_2$) fibrils, the additional density is localized to the C-terminal ends of molecules in the two central subunits. For RT-Aβ40($C_1$) fibrils, the additional density is asymmetric, localized to the C-terminal end of molecules in the upper subunit and to residues 16-20 of molecules in the lower subunit. For RT-Aβ40($C_1$)′ fibrils, multiple layers of strong additional density are apparent.

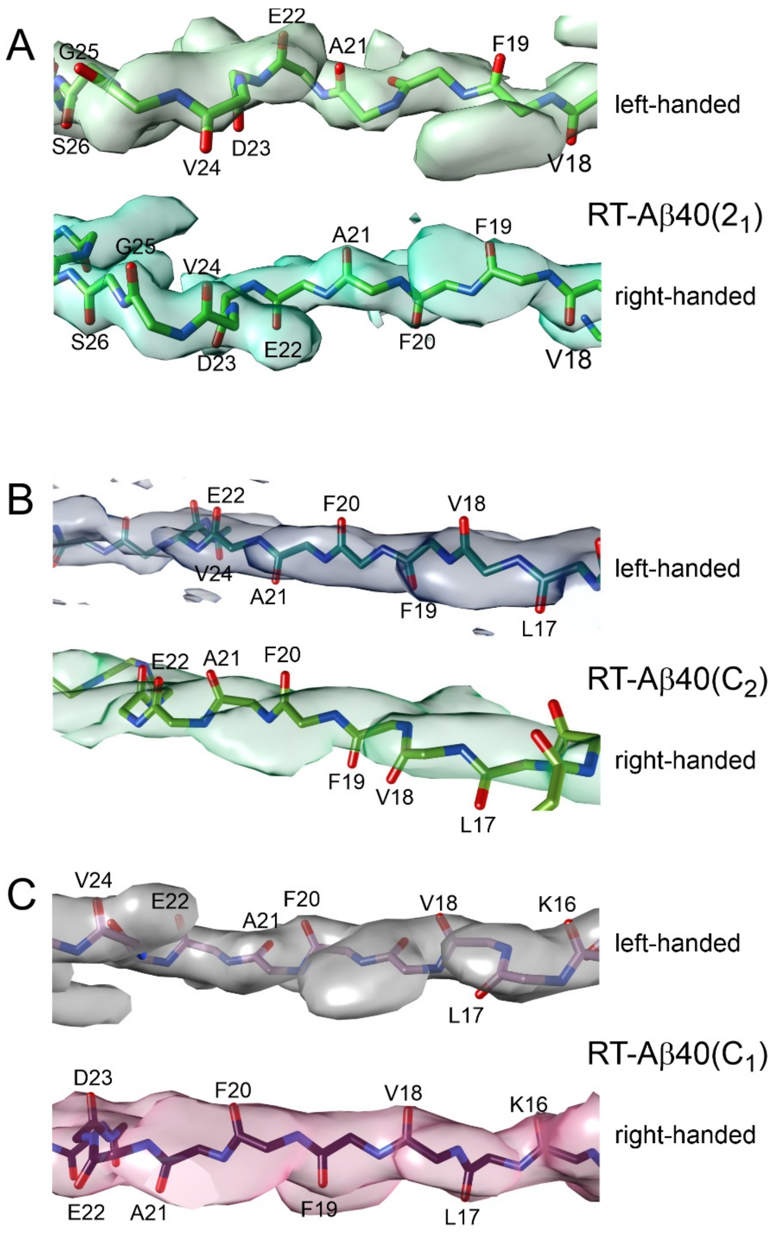


Figure S8: Handedness of helical twist in rapidly twisting Aβ40 fibril polymorphs. (A) Cryo-EM density maps with left-handed and right-handed twist and corresponding molecular models for residues 18-26 in RT-Aβ40($2_1$) fibrils. Backbone carbonyl oxygens align with bulges in the right-handed density, but not the left-handed density. (B) Cryo-EM density maps with left-handed and right-handed twist and corresponding molecular models for residues 19-23 in RT-Aβ40($C_2$) fibrils. Backbone carbonyl oxygens align with bulges in the left-handed density, but not the right-handed density. (C) Cryo-EM density maps with left-handed and right-handed twist and corresponding molecular models for residues 16-24 in the lower subunit of RT-Aβ40($C_1$) fibrils. Backbone carbonyl oxygens align with bulges in the right-handed density, but not the left-handed density. (Backbone conformations in left-handed and right-handed molecular models are significantly different in panels A, B, and C because the molecular models were calculated independently in Xplor-NIH, using either left-handed or right-handed density maps.)

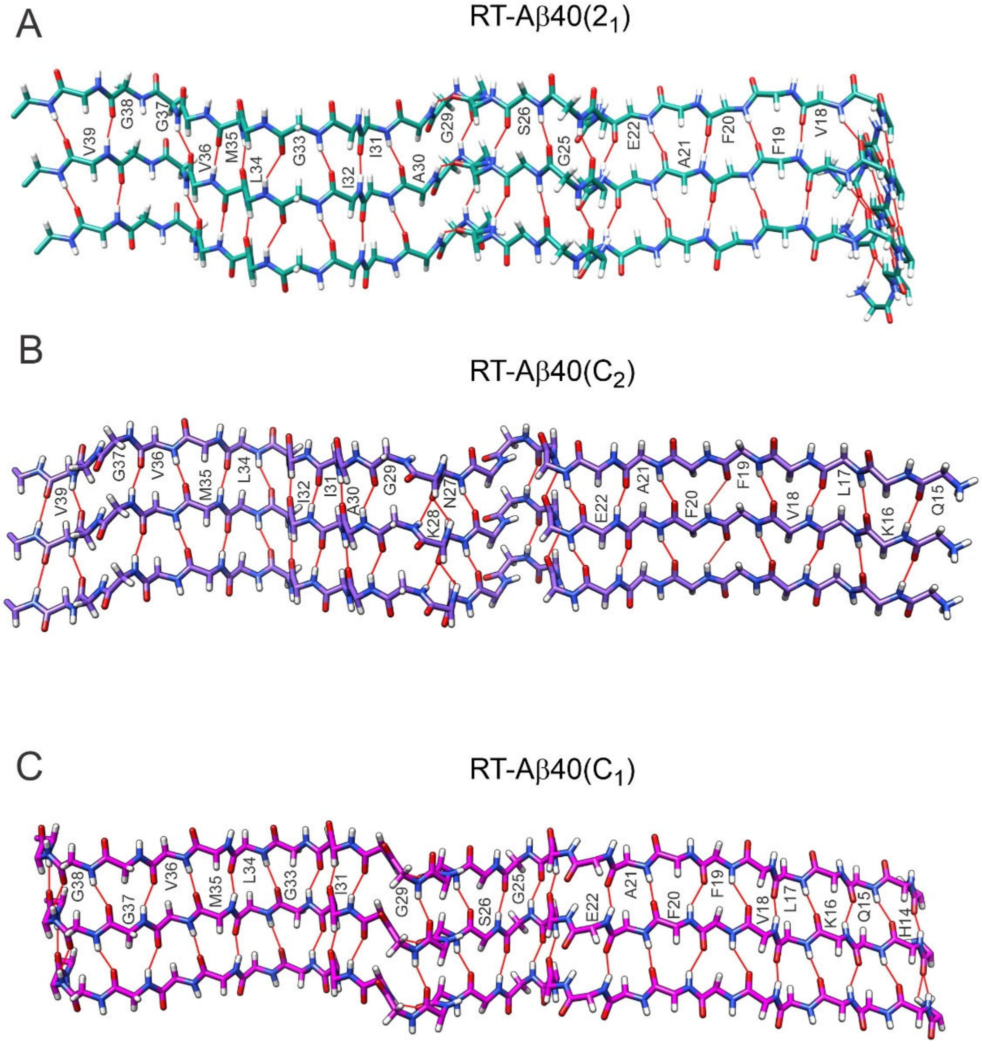


Figure S9: Intermolecular backbone hydrogen bonds in rapidly twisting Aβ40 fibril polymorphs. (A) Three Aβ40 chains from one cross-β subunit of an RT-Aβ40($2_1$) fibrils, viewed perpendicular to main β-sheet plane. Red lines represent hydrogen bonds between backbone carbonyl oxygens of residue i and backbone amide nitrogens of residue i+1. Hydrogen bonds were identified by the criteria of Chimera software, with constraints relaxed by 0.4 Å and 20°. (B) Same as panel A, but for an RT-Aβ40($C_2$) fibril. (C) Same as panel A, but for the lower subunit of an RT-Aβ40($C_1$) fibril.

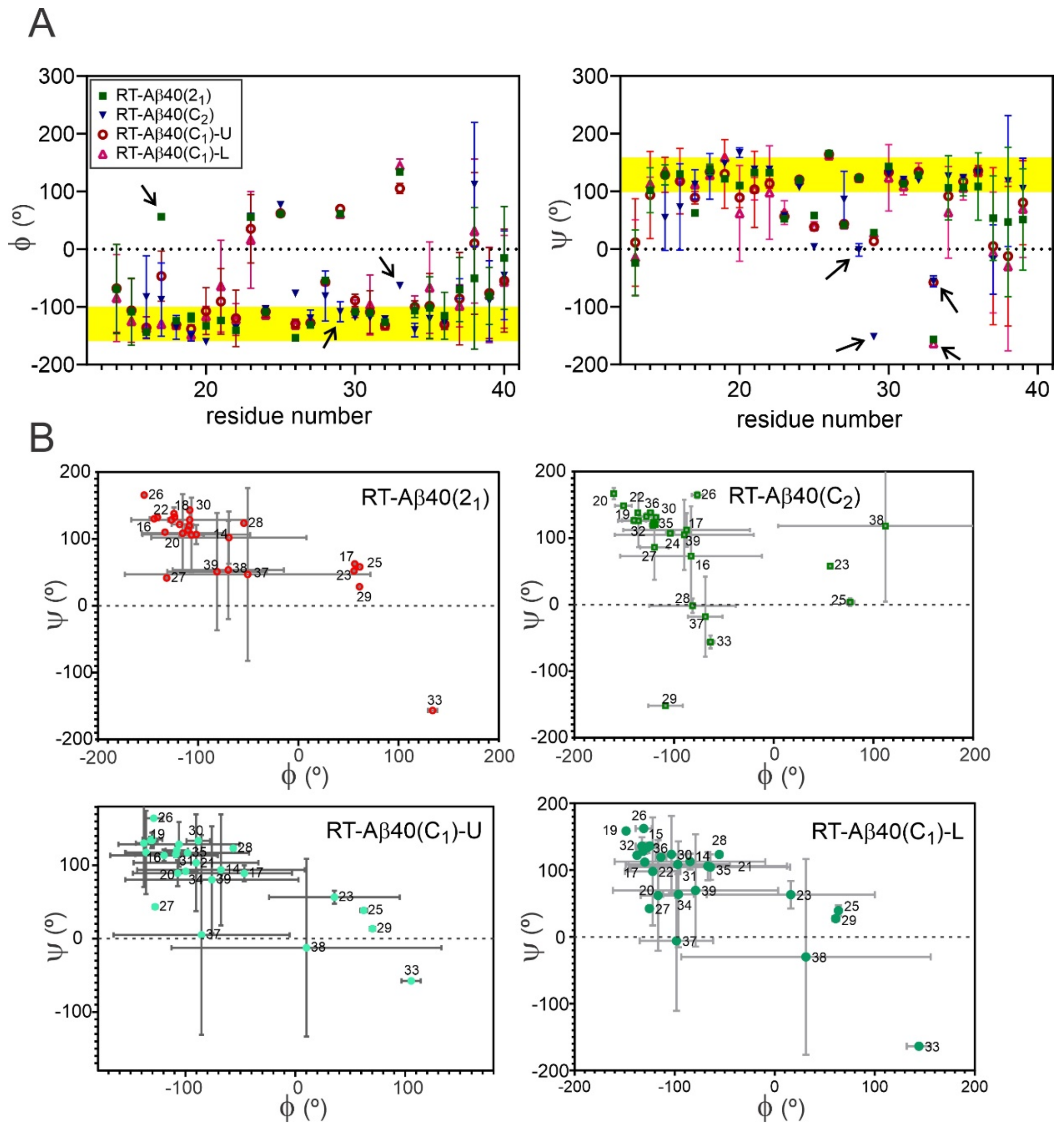


Figure S10: Backbone conformations in rapidly twisting Aβ40 fibril polymorphs. (A) Backbone φ and ψ torsion angles in RT-Aβ40($2_1$), RT-Aβ40($C_2$), and RT-Aβ40($C_1$) fibrils, plotted versus residue number. "U" and "L" represent upper and lower subunits of RT-Aβ40($C_1$) fibrils. Torsion angle values are averages over all molecules within the bundles in Fig. S5, with standard deviations indicated by error bars. Yellow bands indicate values that are typical of β-sheets in proteins. Arrows indicate positions where large conformational differences occur. Large error bars in residues 13-16 and 37-39 are due to the low resolution of the cryo-EM density maps in those regions. (B) Backbone torsion angles represented by Ramachandran plots.

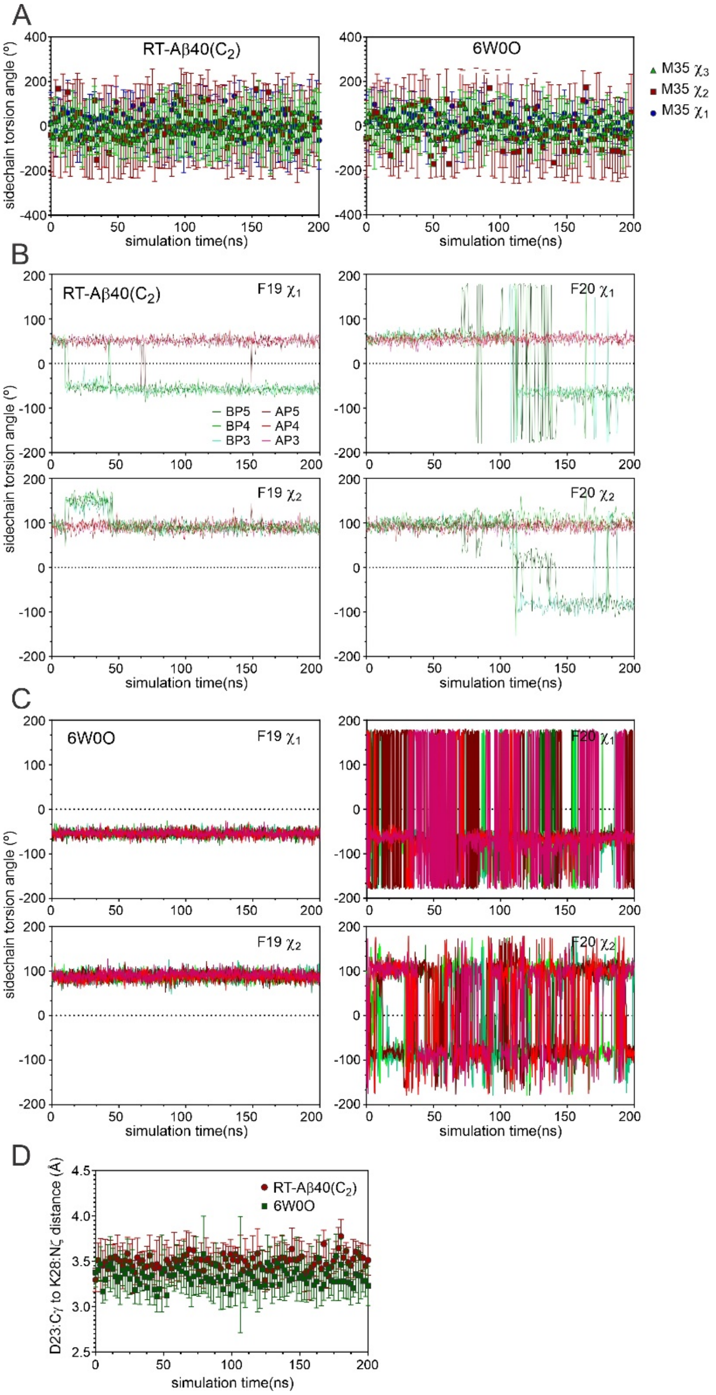


Figure S11: Sidechain dynamics and D23-K28 salt bridges in MD simulations. (A) Time dependences of sidechain $\chi_1$, $\chi_2$, and $\chi_3$ torsion angles of M35 in MD simulations of a rapidly twisting RT-Aβ40(C$_2$) fibril and a slowly twisting brain-seeded fibril (PDB code 6W0O). The all-atom simulations in explicit water solvent with 100 mM NaCl include 14 Aβ40 molecules, or seven β-sheet repeats in each cross-β subunit. Torsion angle values at each time point are averaged over the three central molecules in each subunit, with standard deviations indicated by error bars. Large fluctuations and standard deviations reflect dynamic disorder of M35 sidechains, which are exposed to solvent in both fibril polymorphs. (B) Time dependences of $\chi_1$ and $\chi_2$ torsion angles of F19 and F20 in the rapidly twisting fibril. Values for individual molecules are plotted separately, with AP3, AP4, and AP5 being the central molecules in one subunit and BP3, BP4, and BP5 being the central molecules in the other subunit. F19 sidechains exhibit static disorder, with restricted motions but variable $\chi_1$ values. F20 sidechains are relatively immobile for the first 70 ns, then become partially disordered and dynamic in one subunit. (C) Same as panel B, but for the slowly twisting fibril. F19 sidechains are relatively immobile and conformationally ordered. F20 sidechains are dynamically disordered. (D) Time dependences of distances between sidechain carboxyl carbons of D23 and sidechain amino nitrogens of K28. Values near 3.4 ± 0.3 Å indicate stable salt bridge interactions.

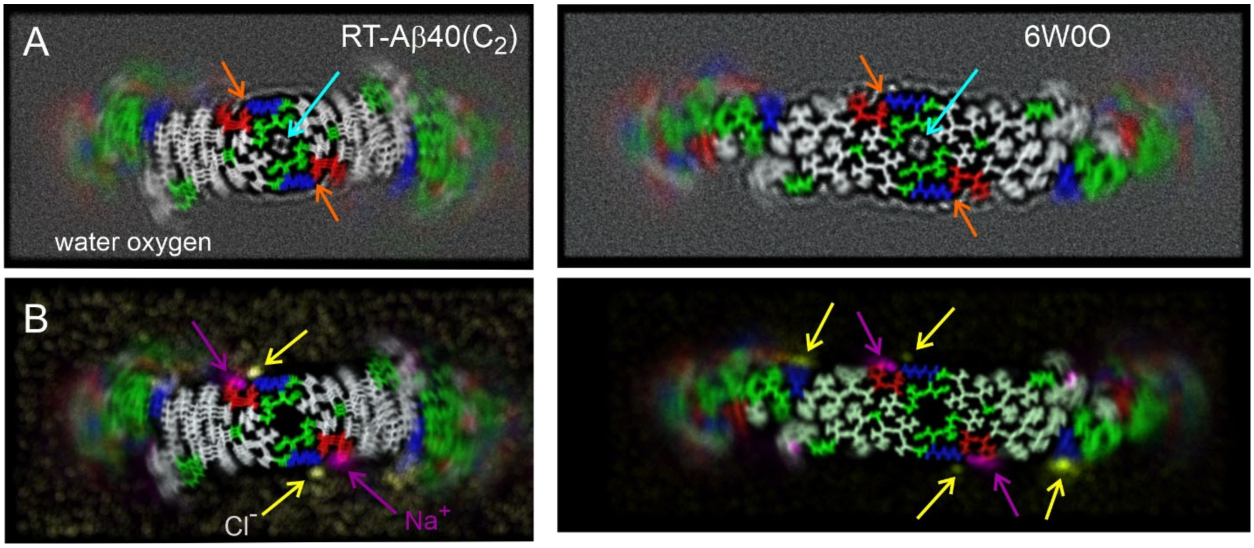


Figure S12: Water and ion distributions from MD simulations. (A) Superpositions of frames from MD simulations of a RT-Aβ40($C_2$) fibril (left) or a fibril with the 6W0O structure (right) in water with 100 mM NaCl. In both cases, 375 frames were combined, covering simulation times from 50 s to 200 ns. Aβ40 molecules are shown in a stick representation, with hydrophobic, polar (and glycine), basic, and acidic residues in white, green, blue, and red, respectively. Oxygen atoms of water molecules are represented by white dots. Water is excluded from both fibril cores, except for the central channels indicated by cyan arrows. Orange arrows indicate stable D23-K28 salt bridge interactions. (B) Superpositions showing the distributions of $Na^+$ and $Cl^-$ ions around the fibrils (magenta and yellow dots and arrows).
